# A screening pipeline identifies a broad-spectrum inhibitor of bacterial AB toxins with cross protection against influenza A virus H1N1 and SARS-CoV-2

**DOI:** 10.1101/2021.08.13.454991

**Authors:** Yu Wu, Nassim Mahtal, Léa Swistak, Sara Sagadiev, Mridu Acharya, Caroline Demeret, Sylvie van der Werf, Florence Guivel-Benhassine, Olivier Schwartz, Serena Petracchini, Amel Mettouchi, Eléa Paillares, Lucie Caramelle, Pierre Couvineau, Robert Thai, Peggy Barbe, Mathilde Keck, Priscille Brodin, Arnaud Machelart, Valentin Sencio, François Trottein, Martin Sachse, Gaëtan Chicanne, Bernard Payrastre, Florian Ville, Victor Kreis, Michel-Robert Popoff, Ludger Johannes, Jean-Christophe Cintrat, Julien Barbier, Daniel Gillet, Emmanuel Lemichez

## Abstract

A challenge for the development of host-targeted anti-infectives against a large spectrum of AB-like toxin-producing bacteria encompasses the identification of chemical compounds corrupting toxin transport through both endolysosomal and retrograde pathways. Here, we performed a high-throughput screening of small chemical compounds blocking active Rac1 proteasomal degradation triggered by the Cytotoxic Necrotizing Factor-1 (CNF1) toxin, followed by orthogonal screens against two AB toxins hijacking defined endolysosomal (Diphtheria toxin) or retrograde (Shiga-like toxin 1) pathways to intoxicate cells. This led to the identification of the molecule N-(3,3-diphenylpropyl)-1-propyl-4-piperidinamine, referred to as C910. This compound induces the swelling of EEA1-positive early endosomes, in absence of PIKfyve kinase inhibition, and disturbs the trafficking of CNF1 and the B-subunit of Shiga toxin along the endolysosomal or retrograde pathways, respectively. Together, we show that C910 protects cells against 8 bacterial AB toxins including large clostridial glucosylating toxins from *Clostridium difficile*. Of interest, C910 also reduced viral infection *in vitro* including influenza A virus subtype H1N1 and SARS-CoV-2. Moreover, parenteral administration of C910 to the mice resulted in its accumulation in lung tissues and reduced lethal influenza infection.

## INTRODUCTION

Host-directed anti-infective chemical compounds that interfere with intracellular membrane trafficking hold promise to alleviate pathophysiological manifestations triggered by bacterial toxins, thus shifting the host-pathogen balance in favor of the host (1). Furthermore, conservation of physicochemical requirements for cell invasion by pathogens and toxins allows to re-direct small molecules targeting endocytic pathways against a wider number of infectious agents notably viruses (2–4). Discovery of such molecules is a current challenge to expand available therapeutics against highly prevalent infections, multidrug-resistant pathogens and to halt pandemics (5).

Bacterial AB toxins are sophisticated nanomachines that rely on both specific physicochemical conditions found in intracellular compartments and accessory cell factors to translocate their enzymatic A-domain into the cytosol. Selective disruption of vesicular trafficking can be leveraged to develop host-directed prophylactic and therapeutic measures against toxin-producing bacteria with broader applications in infectiology (2, 4). To enter the host cells, AB toxins exploit different endocytic pathways which converge towards early endosomes from which they are dispatched to different vesicular pathways (6–8). AB toxins translocate their enzymatic part along the endolysosomal pathway or after retrograde transport from the endoplasmic reticulum (ER), which represents another hot spot of translocation (9, 10). Early endosomes receive cargos, receptors and pathogenic agents coming from the plasma membrane (11). They serve as an essential sorting station to re-route receptors and/or ligands to the cell surface through recycling endosomes, to the trans-Golgi network and the ER through retrograde transport, or the lysosomes for degradation. Each critical step along these vesicular pathways is specified by specialized membrane-associated protein complexes (12). Owing to their initial sorting functions, early endosomes represent targets of interest to develop anti-infectives displaying a large spectrum of action against toxin-producing bacteria and viruses.

The Cytotoxic Necrotizing Factor-1 (CNF1) toxin is produced by extraintestinal pathogenic *Escherichia coli* responsible for urinary tract infections, neonatal meningitis and sepsis (13, 14). The *cnf1* gene also displays a prevalence estimated to 15% in *E. coli* sequence type (ST)131 that underwent an unprecedented global expansion in the last decade and represents a predominant multidrug-resistant (MDR) lineage of *E. coli* in extra-intestinal infections (15). CNF1 binds to Lu/BCAM and 67-kDa laminin receptor (67LR) to enter cells and reaches acidic endosomes from which it translocates into the cytosol to deamidate host Rho GTPases (16–19). The activation of Rho GTPases by CNF1 confers to pathogenic *E. coli* high capacities of host cell invasion (20, 21).

Chemical compounds interfering with the intracellular trafficking of bacterial toxins have shown their efficacy against several infections by bacteria, viruses and parasites *in vitro* and *in vivo* as summarized in Supplementary Table 1 and 2. Known inhibitors act specifically on either of the two main endocytic pathways hijacked by toxins. Retro-2 acts specifically on the endoplasmic reticulum (ER) exit site component Sec16A thereby compromising the interaction between the retrograde trafficking chaperone GPP130 and syntaxin-5 for proper transport of Shiga toxin to the ER (22, 23). Both inhibitors EGA and ABMA affect exclusively the trafficking of toxins along the endolysosomal pathway by unidentified mechanisms (2, 4). Currently, no single compound conferring protection to AB toxins hijacking either the endolysosomal or retrograde pathways are available.

Here, we developed a high-throughput screening (HTS) pipeline allowing the identification of a small chemical compound that protects host cells against eight unrelated bacterial AB toxins. Moreover, C910 properties can be leveraged to confer protection of mice against flu caused by influenza A virus subtype H1N1 and to reduce cell infection by SARS-CoV-2. Mechanistically, we provide evidence that C910 interferes with the sorting function of EEA1/Rab5-positive early endosomes.

## RESULTS

### Screening of compounds conferring resistance to a large spectrum of AB toxins

We developed a pipeline aiming at screening small chemical compounds protecting host cells against a broad spectrum of bacterial AB toxins hijacking either the endolysosomal (CNF1 and diphtheria toxin, DT) or the retrograde pathways (Shiga-like toxin 1, Stx1) (Figure 1A). We first performed HTS of 16,480 available chemical compounds from the ChemBridge DIVERSet library for their capacity to protect primary human umbilical vein endothelial cells (HUVECs) against CNF1-induced cellular depletion of Rac1 (Figure 1B). The CNF1 toxin hijacks the endolysosomal pathway to penetrate into cells and catalyzes the deamidation of a specific glutamine residue on small Rho GTPases, notably Rac1 (17, 18). This post-translational modification activates Rac1 and thereby sensitizes this GTPase to ubiquitin-mediated proteasomal degradation (24). Contrary to previous screens for antitoxins based on terminal cytotoxic effects, the readout of the screen presented here relies on direct monitoring of the target of CNF1. Briefly, cells were treated with the CNF1 toxin at 10 nM to achieve maximal proteasomal degradation of Rac1 in 6 h. Quantitative immunofluorescence staining of Rac1 cellular levels was applied directly to the cells. The threshold of the chemically-induced protection against Rac1 depletion was set to 30% to sort robust candidate hits (Figure 1B).

**Figure 1.**
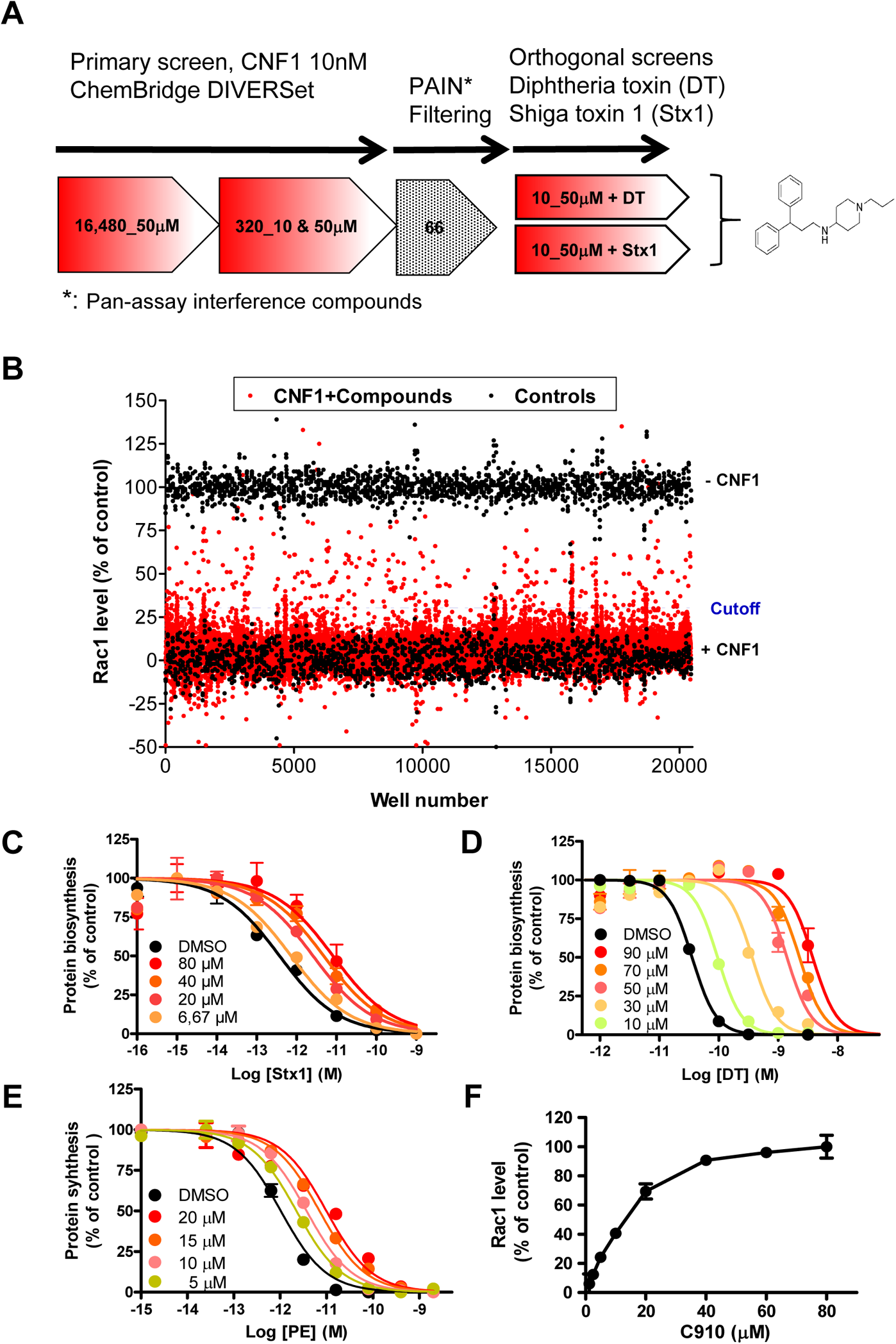
High-throughput screen of broad-spectrum anti-toxin inhibitors. A: General scheme and results of the screening pipeline developed to isolate small chemical compounds inhibiting CNF1 and bacterial AB toxins hijacking the endolysosomal (DT for diphtheria toxin) or retrograde (Stx 1 for Shiga like toxin) pathways, respectively. Chemical structure of N-(3,3-diphenylpropyl)-1-propyl-4-piperidinamine, referred to as C910. B: ChemBridge library compounds were screened on HUVECs. Upper black points represent positive controls (100% Rac1 signal in control HUVECs), lower black points are negative controls corresponding to Rac1 level set to 0% in HUVECs exposed to 10 nM CNF1, 6 h, while red points correspond to HUVECs incubated with library compounds in the presence of 10 nM CNF1, 6 h. Cutoff was set for 30% of Rac1 signal. C-E: Graphs show protein synthesis inhibition induced by Stx1, DT and PE. (C and D) HeLa cells were pretreated 1 h in DMEM with C910 at indicated concentrations (color filled circles), or DMSO vehicle (black circles) prior to addition of increasing concentrations of Stx1 (C) or DT (D) for 18 h and 6 h, respectively. Each point represents the mean of duplicates ± s.e.m. of one representative experiment (*n* = 2). E: L929 cells were incubated for 1 h with C910 (color filled circles), or with DMSO vehicle (black circles) before addition of PE for 18 h at indicated concentrations. Protein synthesis was measured by scintillation counting. Each point represents the mean of duplicates ± s.e.m. of one representative experiment (*n* = 2). F: Graph shows Rac1 cellular level in the presence of increasing concentrations of C910. HUVECs were intoxicated with 10 nM CNF1 for 6 h in the presence or absence of C910 at the indicated concentrations. Each point represents the mean of duplicates ± s.e.m. of one representative experiment (*n* = 2).

In total, 320 hits were cherry-picked and subjected to a second round of screening at two concentrations in triplicate. 66 compounds passed the second round of screening. These hits were subsequently reordered and freshly prepared for further validation and then filtered from pan-assay interference compounds (PAINS), i.e. unstable molecules, irreversible modifiers or compounds that are frequently active in other screens (25). A total of 10 compounds were shortlisted as robust inhibitors of CNF1-mediated cellular degradation of Rac1. These compounds were subsequently analyzed for their protective action against two bacterial toxins following either endolysosomal or retrograde pathways. These orthogonal screens were set to counter-select compounds acting directly on the proteasomal degradation of Rac1, and to identify compounds endowed with a large spectrum of action against AB toxins. Here, we made use of diphtheria toxin (DT), as it shares with many enzyme-based AB toxins the need to reach early acidic compartments along the endolysosomal pathway from which the enzymatic A-domain translocates into the cytosol (26, 27). The ADP-ribosyltransferase activity of DT poisons host cells by catalyzing the post-translational modification of elongation factor-2 (EF-2), thereby blocking protein synthesis. We also made use of Stx1 as it undergoes retrograde transport from the early endosomes to the Golgi before reaching the ER where it exits into the cytosol (10, 28). The *N*-adenine glycohydrolase activity of Stx1 inhibits protein biosynthesis. The cytotoxic effects induced by DT and Stx1 were quantified by a measure of [^14^C]-leucine incorporation into neosynthesized proteins. Among the 10 compounds tested, three molecules turned to be active against DT and one of these was active against Stx1 as well. Taken collectively, this screening strategy allowed us to isolate one compound out of a total of 16,480 molecules, here referred to as C910. The chemical structure of N-(3,3-diphenylpropyl)-1-propyl-4-piperidinamine can be divided into four parts (Figure 1A). The western half bears two aromatic phenyl groups while the eastern part bears a central amino group, a piperidine ring with a terminal propyl chain.

We next conducted a dose-response analysis to determine the relationship between compound concentration and C910 antitoxin activity. We determined half maximal effective concentrations (EC_50s_) of 35.6 μM for Stx1 and 60.1 μM for DT, as well as 4.9 μM for Stx2 (Figure 1C-D and Sup. Figure 1). We also evaluated the effective concentration of C910 against Exotoxin A (PE) from *Pseudomonas aeruginosa*, that shares with DT and STx the capacity to inhibit protein biosynthesis. This toxin exploits intertwined endolysosomal and retrograde pathways (29, 30) to reach the ER and get extruded into the cytosol (7, 31). Cells were intoxicated with increasing doses of PE in the presence of C910 or DMSO vehicle establishing a value of EC_50_ of 12.6 μM (Figure 1E and Sup. Figure 1A-D). Finally, we determined that C910 inhibited CNF1 (10 nM) with an IC_50_ = 11.3 μM (Figure 1F). We showed that C910 exhibits no detectable toxicity on HUVECs and HeLa cells at concentrations below 75 μM (Sup. Figure 1E). Thus, we have identified a chemical entity that decreases the susceptibility of host cells to five different bacterial toxins exploiting defined intracellular trafficking pathways of intoxication.

### C910 inhibits CNF1-mediated invasion of cells by UPEC

Invasion of epithelial and endothelial cells by UPEC leads to severe and recurrent forms of infections (32). Here, we investigated whether C910-induced resistance of cells to CNF1 cytotoxicity could decrease the susceptibility of host cells to bacterial invasion. The protective action of C910 was assessed in gentamicin-protection experiments, upon infecting cells with a *cnf1*-deleted strain of UPEC in the control condition versus complementation with exogenously added CNF1 at 1 nM. This setting allowed to measure reproducible rates of cell invasion (33). Remarkably, cells treated with C910 at 15 μM were protected against CNF1-mediated invasion by UPEC without affecting bacterial adhesion to the cells (Figure 2A and 2B). C910 had no impact on bacterial viability (data not shown). We also verified in parallel that C910 abolished CNF1-mediated depletion of Rac1 during cell infection (Figure 2C). In parallel, we found no protective effect of C910 on the Cdc42/Rac1-dependent invasion of cells by *Salmonella typhimurium* (Figure 2D and E). In contrast to CNF1 toxin-producing UPEC, cell-bound *S. typhimurium* directly injects Cdc42/Rac1-activating effectors through the type-3 secretion system (T3SS)-1, thereby bypassing endocytosis and intracellular trafficking steps required for AB toxins action (34). The specificity of chemically-induced cellular protection against UPEC points toward an action of C910 on the endocytic machinery that is exploited by CNF1, rather than an action on Cdc42/Rac1-driven large-scale membrane deformations that are at play for cell invasion by pathogenic bacteria.

**Figure 2.**
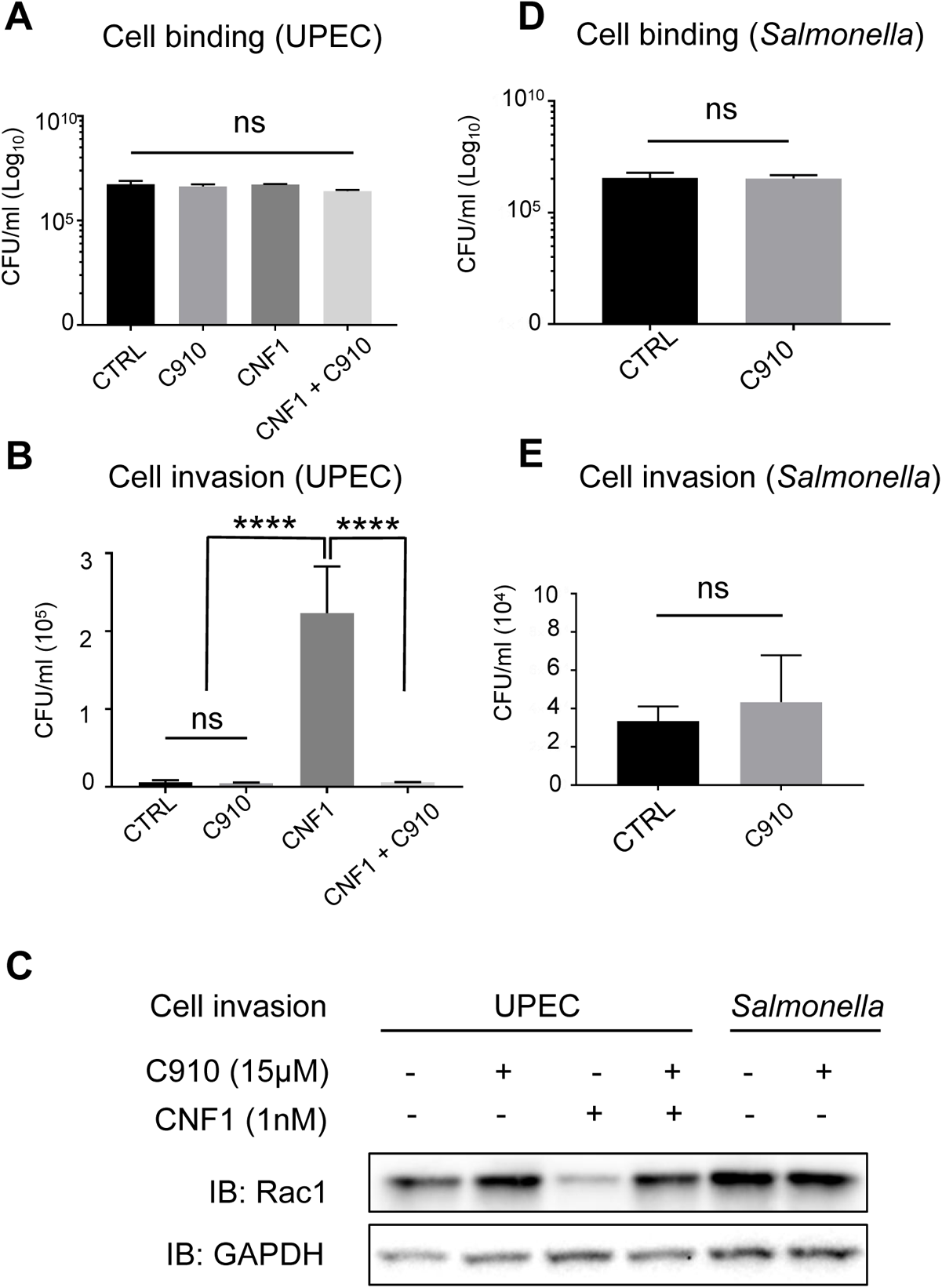
C910 protects host cells against UPEC invasion mediated by CNF1. A-E: Measure of cell invasion by uropathogenic *E. coli* UTI89Δ*hlyA*Δ*cnf1* (UPEC) and *Salmonella enterica* serovar Thyphimurium SL1344 (Salmonella). HUVECs were pre-incubated 30 min with 15 μM C910 or vehicle prior to infection during 2.5 h by UPEC in the absence or presence of 1 nM CNF1. HUVECs were incubated 3 h with 15 μM C910 or vehicle prior to infection with *Salmonella* in the presence or absence of C910 during 1 h. Graphs show number of viable bacteria associated to the cells (A and D) or intracellular bacteria resistant to gentamicin treatment (B and E). Values correspond to colony forming units (CFU)/ml, presented as mean of ± s.e.m. from three independent experiments. One-way ANOVA was performed overall (p < 0.0001) with Tukey’s multiple comparison test to compare the treated conditions in (A) and (B), **** *p* < 0.0001 or ns: not significant. Data in and (E) were analyzed with unpaired two-tailed *t*-test, ns: not significant. (C) Immunoblot anti-Rac1 shows that C910 blocked CNF1-mediated Rac1 depletion during cell infection by UPEC, as described in (B) and that C910 and/or *Salmonella* infection, as described in (E), did not affect Rac1 cellular levels. Anti-GAPDH immunoblot was used as a loading control.

### Extended spectrum of protection against bacterial AB-like toxins

We pursued the characterization of the antitoxin spectrum of C910 by studying a paradigm of AB toxins that translocate their A-enzymatic components at acidic pH through a pore formed by B-subunit oligomers. Anthrax toxin from *Bacillus anthracis* is composed of the protective antigen (PA), lethal factor (LF) and edema factor (EF) (35). The 63-kDa matured form of PA undergoes octamerization at the cell surface in association with receptors leading to toxin-receptor endocytosis (36–38). The acidic environment of late and matured early endosomes then drives the insertion of oligomerized PA into the lipid bilayer through which LF and/or EF translocate into the cytosol either directly or via a two-step mechanism involving internal vesicles of late endosomes (39). HUVECs were intoxicated with the lethal toxin (LT: PA + LF) at two different concentrations sufficient to induce a complete cleavage of MEK2 substrate of LF, between 2 and 4 h (Figure 3A). We found that addition of C910 at 40 μM markedly delayed the kinetics of MEK2 cleavage by protection factors comprised between 2.2 (LT) < P < 2.5 (LT_1/10_) (Figure 3A and 3B). This demonstrates the protection conferred overtime by C910 to host cells intoxicated by LT from *B. anthracis*.

**Figure 3.**
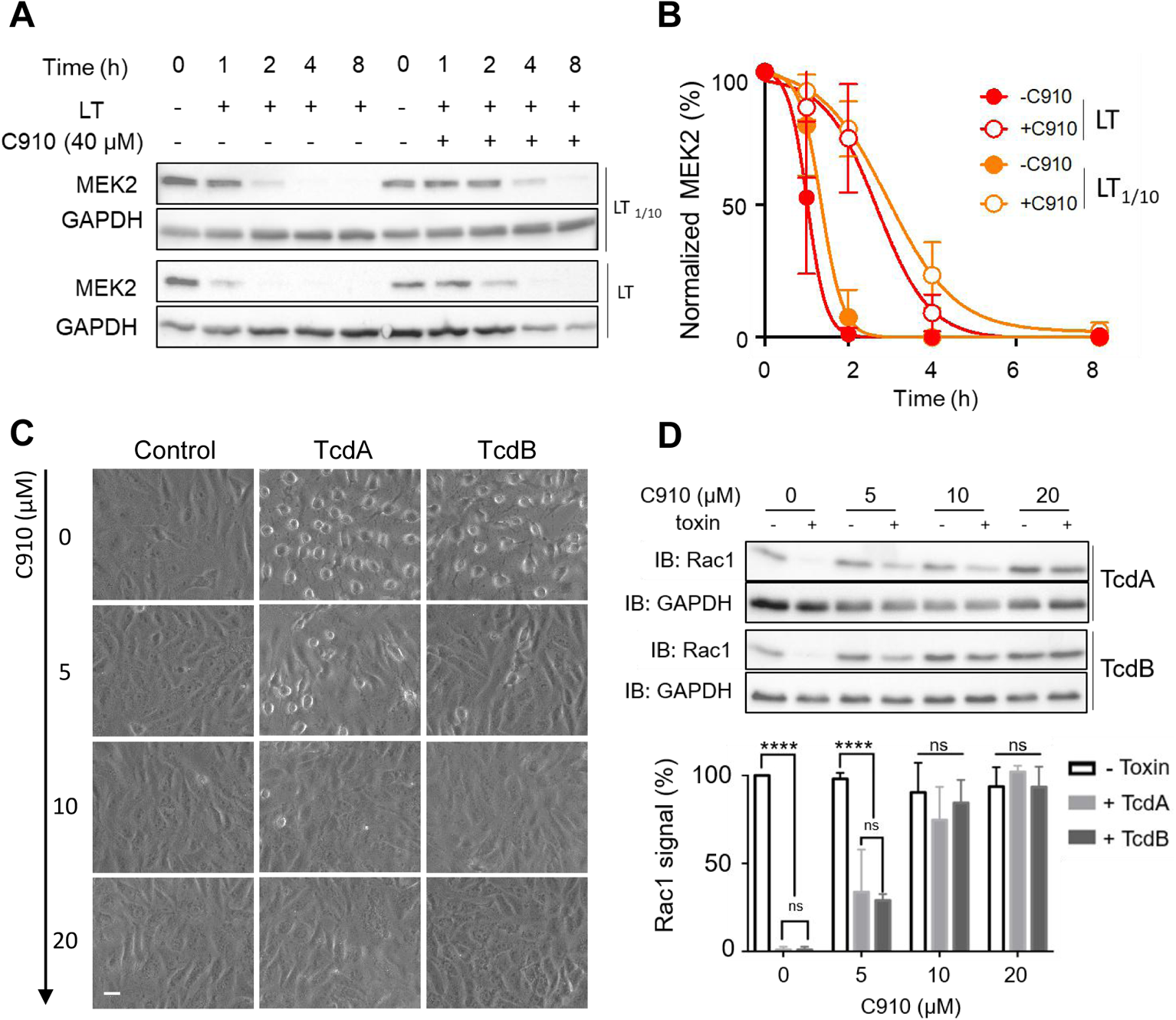
C910 protects host cells against lethal toxin from *B. anthracis* and large glucosylating toxins from *C. difficile.* A: Immunoblots show kinetics of MEK2 cleavage at two concentrations of lethal toxin, PA 0.3 μg/ml / LF 0.1 μg/ml (LT) or diluted 1/10 (LT_1/10_) together with the protection conferred by C910 at 40 μM. Anti-GAPDH immunoblots show loading control. The blots are representative of four independent experiments. B: Graph shows nonlinear regression curves of normalized MEK2 level as a function of time (h) (R^2^ > 0.9), in conditions described in (A). Percentage of MEK2 signal in HUVECs intoxicated for the indicated periods of time with LT (red circles) and LT_1/10_ (orange circles) in the presence of DMSO vehicle (plain circles) or C910 at 40 μM (open circles). Each point represents mean ± s.d. from four independent experiments. Data were fitted to obtain the mean time (t_1/2_) of MEK2 cleavage and the protection factors (i.e., P = t_1/2, LT+C910_ / t_1/2, LT_). C: Vero cells intoxicated with 8 cytotoxic units (cu) of TcdA or TcdB for 8 hours in the presence of DMSO vehicle or C910 at indicated concentrations. Scale bar, 10 μm. D: Anti-Rac1 immunoblots show the non-glucosylated form of Rac1. Vero cells were treated as described in (C). Lower graph corresponds to Rac1 signals normalized to GAPDH and set to 100% for control condition, data are expressed as mean ± s.d. from three independent experiments. Two-way ANOVA with Tukey’s multiple comparison test was used to compare toxin-treated cells with non-intoxicated controls in the presence of C910 at different concentrations, **** *p* < 0.0001 or ns: not significant.

We went on to test the effect of C910 on large clostridial glucosylating toxins (LCGTs) TcdA and TcdB from *Clostridium difficile* (40). These toxins hijack different host receptors to enter into cells via endocytosis and translocate from acidified endosomes (41–43). The internalization of LCGTs occurs via a dynamin-dependent process that largely involves clathrin. LCGTs glucosylate a threonine residue in the effectors-binding loop of Rho GTPases to short circuit the signaling cascades that control the actin cytoskeleton. Consequently, this produces a rounding of cells. Highly sensitive Vero cells were intoxicated with 8 cytotoxic units of TcdA or TcdB. These conditions were set to reach 100% of cell rounding at 8 h of intoxication. In these conditions, co-treatment of cells with C910 at 5 μM markedly reduced the percentage of intoxicated cells (Figure 3C). The extent of protection conferred by C910 on LCGTs cytotoxicity was quantified by anti-Rac1 immunoblotting (Figure 3D). This is based on the observation that glucosylation of the threonine-35 of Rac1 blocks its recognition by the anti-Rac1 [clone 102]-monoclonal antibody (44). This allowed direct visualization of the dose-dependent inhibition conferred by C910 on LCGTs-driven post-translational modification of Rac1, which was maximal at 20 μM (Figure 3D).

Altogether, our results show that C910 confers to host cell resistance against eight bacterial AB toxins relevant to public health or produced by a pathogen of concern of misuse.

### C910 acts between CNF1 cell entry and catalytic steps

We showed that C910 protects cells against multiple AB toxins with distinctive catalytic activities and host cell receptors, thereby pointing for a direct action on host cells. Nevertheless, we verified the absence of an inhibitory effect of C910 on CNF1-mediated deamidation of RhoA measured *in vitro* (Sup. Figure 2A). We also verified that C910 did not affect the association of CNF1 to cell surface-exposed receptors. For this, CNF1 was chemically coupled to the Cy3 fluorophore (CNF1-Cy3) and incubated with cells at 4°C to allow its binding to host cells. Fluorescence-activated cell sorting (FACS) analysis showed that a concentration of C910 above its IC_50_ against CNF1, did not significantly alter the mean fluorescence intensity (MFI) of CNF1-Cy3 bound to the cells (Figure 4A). We also recorded an absence of effect of C910 on the cellular binding of the B-subunit of Shiga toxin (STxB) (Sup. Figure 2B). Furthermore, C910 had no detectable effect on the internalization of CNF1-Cy3 and STxB (Figure 4B). Taken collectively, these data indicate that C910 operates neither at initial steps of host cell receptor binding and endocytosis of CNF1 and STxB nor at a later stage of CNF1 enzymatic activity.

**Figure 4.**
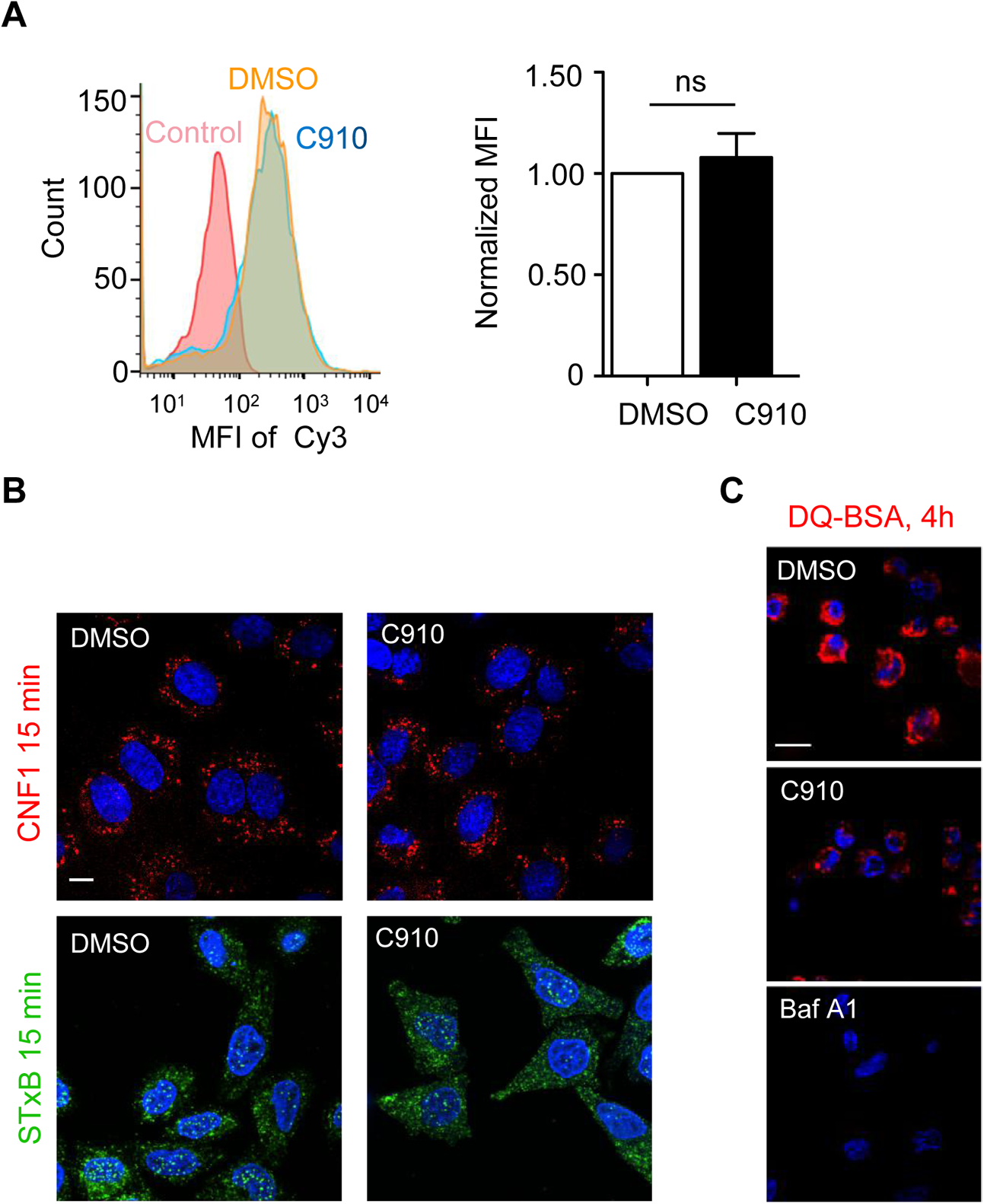
Absence of impact of C910 on CNF1 and STxB entry into cells. A: HUVECs were pre-treated with 40 µM C910 or DMSO for 2 h, detached and incubated with labeled CNF1-Cy3 (1/500) on ice for 30 min in the presence or absence of C910 prior to flow cytometry analysis. Signal of non-treated cells was used as background. MFI (Mean Fluorescence Intensity) of C910-treated cells was normalized to that of DMSO-treated cells. Histogram represents mean ± s.e.m. of three independent experiments. ns: not significant, paired two-tailed *t*-test. B: Representative images show the internalization of CNF1-Cy3 (red) in HUVECs and STxB-his (green) in HeLa cells, nuclei were stained with DAPI (blue). Cells were pre-treated with C910 or DMSO for 30 min at 37°C, then incubated with CNF1-Cy3 (red) or STxB-his (green) on ice for 30 min and followed by incubation for 15 min at 37°C in the continues presence of C910 or DMSO. Scale bar, 20 µm. C: Representative images show the level of DQ-BSA cleavage (red) in live cells treated 4 h with DMSO, 20 µM C910 or 100 nM Bafilomycin A1 (Baf A1). Scale bar, 20 µm.

These exploratory results prompted us to examine a broader impact of C910 on the intracellular transport to lysosomes, by using a protease-dependent dequenching of self-quenched red fluorescent DQ^TM^ Red BSA (DQ-BSA). At a late timing of the chase of 18 h, we observed an accumulation of cleaved DQ-BSA with no detectable effect of C910 (data not shown). When the chase was carried out during a shorter period of 4 h, we observed a strong reduction of cleaved DQ-BSA in C910-treated cells, as compared to control conditions (Figure 4C). In parallel, we recorded no significant effect of C910 on lysosomal cathepsin B activity measured *in vitro* and in cells (Sup. Figure 3A and 3B). Together, these data argue for an action of C910 on endolysosomal trafficking thereby preventing toxin actions.

### C910 affects early endosome homeostasis and sorting of toxins

The above data prompted us to analyze the impact of C910 treatment on the morphology of vesicular compartments along the endolysosomal and retrograde pathways. This was conducted after a short period of time to avoid cascading effects on the endomembrane system. Immunolabeling of several protein markers failed to detect a significant effect of C910 on the distribution and morphology of endolysosomal or retrograde pathway-related compartments except for Rab5-positive early endosomes (Figure 5A). Indeed, compartments positive for Rab5 and the early endosome antigen 1 (EEA1) appeared significantly enlarged and devoid of the late endosomal marker lysobisphosphatidic acid (LBPA) (Figure 5B and Sup. Figure 3C). The extent of swelling of EEA1-positive compartments increased as a function of time (Sup. Figure 4A) and dose of C910 (Sup. Figure 4B). Transmission electron microscopy analysis of BSA-gold loaded endosomes from HUVECs, at an early time point of endocytosis, showed vacuoles containing gold-particles that had an electron-lucent lumen and few internal vesicles (Sup. Figure 4C). In parallel, we recorded an increase in the size distribution of BSA-gold positive compartments (Sup. Figure 4C).

**Figure 5.**
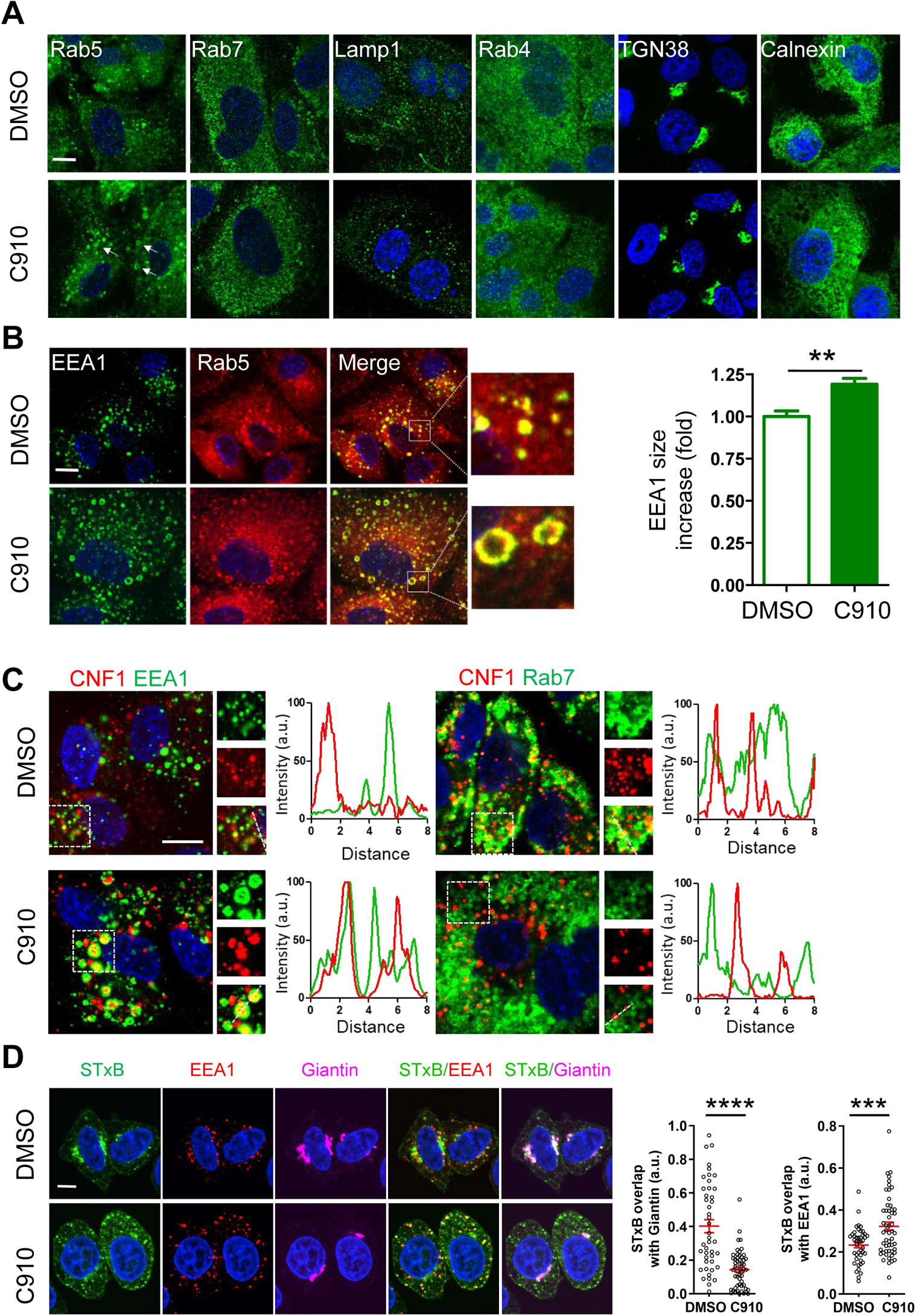
C910 alters EEA1-positive early endosomes morphology and function. A-D: immunofluorescence analyses of C910 specific effect on EEA1/Rab5 compartments and CNF1 or STxB progression along the endolysosomal and retrograde pathways, respectively. Nuclei were labeled with DAPI (blue). Scale bar, 10 µM. A: HUVECs were treated with 40 µM C910 or DMSO for 45 min and labeled for the indicated markers of intracellular compartments. B: Images of EEA1- and Rab5-positive compartments in HUVECs treated as in (A). The size of EEA1-positive particles was quantified and normalized to control, and displayed as mean ± s.e.m., *n* > 20 cells in one representative experiment (*n* = 2). ** *p* < 0.01, unpaired two-tailed *t*-test. C: Images of EEA1- and Rab7-positive compartments in HUVECs pre-treated with 40 µM C910 or DMSO vehicle for 30 min followed by incubation with CNF1-Cy3 for 90 min in the presence of C910 or DMSO. C: Line plots show the intensity of fluorescence along the line. D: HeLa cells were pre-treated for 5 min with 40 µM C910 or DMSO before addition of Alexa Fluor 488-STxB (STxB, 1/5000, 4°C for 30 min; green signal), followed by incubation for 55 min at 37°C in the presence of C910 or DMSO. Cells were then labeled for EEA1 (red) and Giantin (pink). Right graphs show Mander’s overlap coefficients between STxB and EEA1 as well as STxB and Giantin, expressed as arbitrary unit (a.u.). Data show mean ± s.e.m. for 44 (DMSO) and 53 (C910) cells from three independent experiments. **** *p* < 0.0001, unpaired two-tailed *t*-test.

We went on to define how C910 treatment affects the trafficking of CNF1. After 90 min of intoxication, CNF1-Cy3 accumulates in Rab7-positive late endosomes (Figure 5C). In contrast, we observed that CNF1-Cy3 accumulates in the lumen of enlarged EEA1-positive endosomes in C910-treated HUVECs (Figure 5C). This was accompanied with a significant decrease of co-localization between CNF1-Cy3 and Rab7 signals in C910-treated cells compared with control cells (Sup. Figure 5A). In parallel, we found that C910 significantly promotes the accumulation of the Alexa Fluor 488 conjugated STxB in EEA1-positive endosomes at the expense of STxB accumulation in the Golgi area, defined by Giantin immunostaining (Figure 5D).

Altogether, our results establish that C910 acts on EEA1-positive early endosomes homeostasis compromising the progression of two AB toxins along their respective endolysosomal or retrograde pathways.

### C910 inhibits cell infection by influenza A virus H1N1 and SARS-CoV-2

The emergence of SARS-CoV-2, the etiologic agent of coronavirus disease (COVID) 2019, resulted in a global pandemic affecting more than one billion people. Flu pandemics include the devastating Spanish flu (> 50 million death worldwide) and the more recent 2009 H1N1 pandemic. These repeated pandemic episodes of severe pneumonia of viral origin call for developing broad anti-infective strategies against viral pathogens. Though more sophisticated than toxins, several viruses share with them physicochemical and host proteases requirements found inside endosomes to invade host cells. Well-defined influenza A virus (IAV) trafficking in host-cell compartments shows that viral particles enter cells by multiple endocytic mechanisms and traffic through EEA1-positive early endosomes prior to the release of genetic material into the cytosol from late endocytic compartments (45, 46). Infection of cells by SARS-CoV-2 involves its binding to the angiotensin converting enzyme 2 (ACE2) receptor (47) and processing of the spike glycoprotein by host proteases at the plasma membrane and in late endosomes for membrane fusion (48). Processing of the spike protein expressed at the plasma membrane of infected cells by TMPRSS2 serine protease also facilitates cell-cell fusion in syncytia formation (49).

We first examined the effect of C910 on a single IAV infectious cycle. A549 human pulmonary epithelial cells, pretreated with C910 or vehicle alone, were infected with a reporter H1N1_WSN_ virus expressing the mCitrine fluorescent protein. Expression of mCitrine from this reporter virus reflects viral genome replication (50). We monitored the efficiency of cell infection by flow cytometry at a short timing of 6 h and recorded that C910 treatment reduced the percentage of infected cells by 3.6 fold (Figure 6A, left panels). We then monitored the effect of C910 on multiple-cycle of infection. At a longer time of infection, C910 decreased the viral titer by 25.8 fold (Figure 6A, right panel), thereby establishing the strong inhibitory action of C910 on cell infection by IAV.

**Figure 6.**
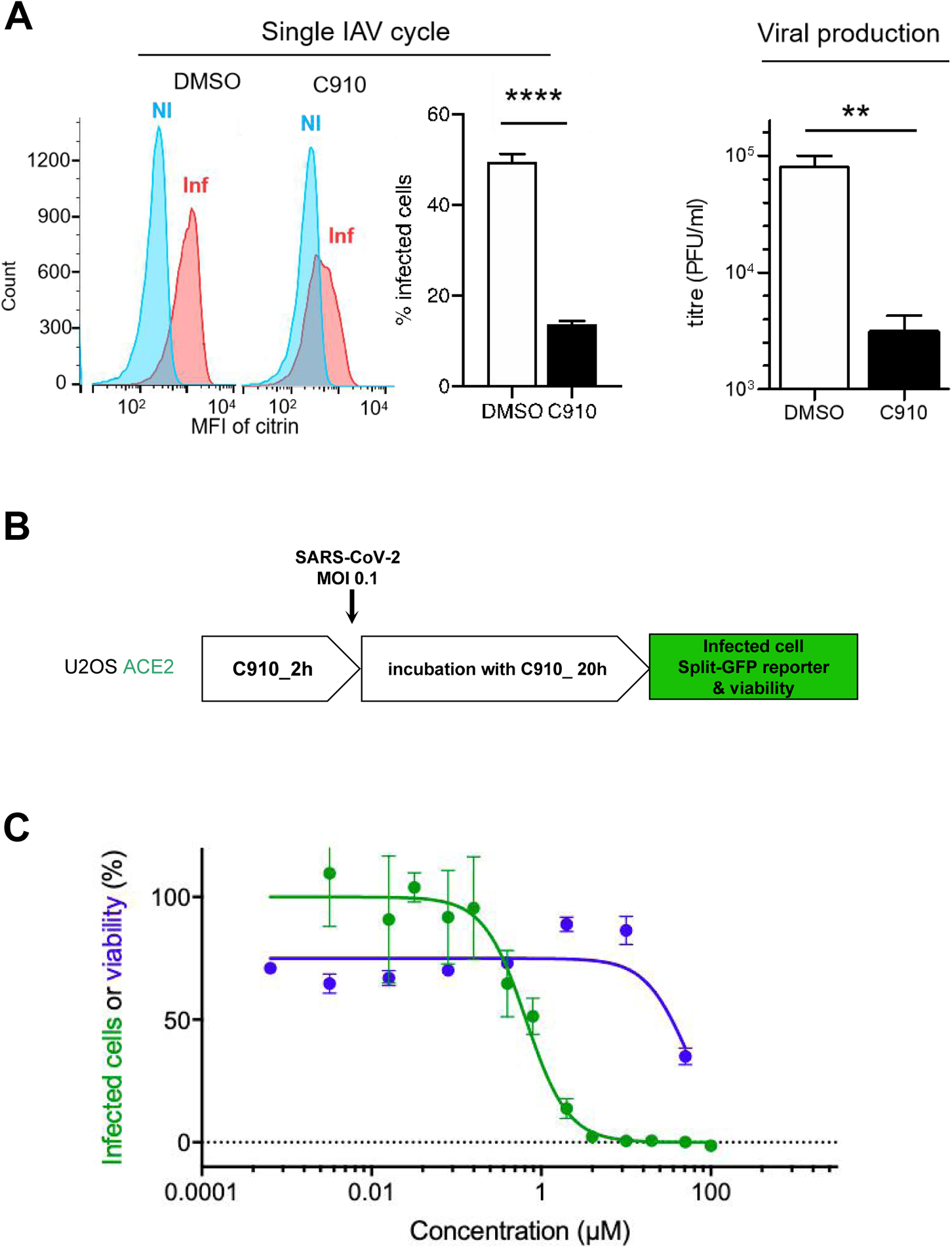
C910 inhibits cellular infection by IAV H1N1 and SARS-CoV-2. A: Left, Effect of C910 on IAV H1N1 single cycle. A549 were pre-treated with C910 (30 µM) or DMSO alone for 1 h at 37°C prior to infection by H1N1_WSN_ mCitrin reporter at a MOI of 5 in the presence of C910 30 µM or DMSO. At 6 h post-infection, cells were analyzed by flow cytometry. Mean Fluorescence Intensity (MFI) of non-infected or infected cells are shown. Histogram represents mean ± s.e.m. of 6 replicates in one representative experiment. **** *p* < 0.0001, unpaired two-tailed *t*-test. Right, Inhibitory effect of C910 on H1N1 viral production. A549 were infected with H1N1_pdm09_ MOI at 0.001 in the presence of 30 µM C910 or DMSO alone. Viral production was titrated 24 h later on cell supernatant by plaque forming assay. Histogram represents mean ± s.e.m of 3 replicates in one representative experiment. ** *p* = 0.005, unpaired two-tailed *t*-test. B: Schematic protocol of SARS-CoV-2 infection. U2OS-ACE2 GFP1-10 and GFP 11 cells were pre-incubated with C910 for 2 h and then SARS-CoV-2 (MOI = 0.1) was added for 20 h. The relative infection efficiency was calculated as the ratio between the GFP-area and the total amount of cells stained with DAPI. C: Data are shown as mean ± s.d. from duplicate wells, one representative experiment (*n* = 3).

As for many viruses, SARS-CoV-2 relies on components of endocytic pathways to infect cells (51, 52). This encompasses the activity of PIKfyve kinase that resides predominantly on early endosomes to regulate endomembrane homeostasis, also being the specific target of Apilimod, a compound that blocks SARS-CoV-2 replication (51). We tested the potential protective effect of C910 on U2OS cells expressing human ACE2 (U2OS-ACE2), which are highly sensitive to SARS-CoV-2 (49). These cells express functional GFP upon productive infection and fusion with neighboring cells, providing a convenient readout of infection. U2OS-ACE2 were preincubated with C910 before viral exposure, and viral infection was revealed after a long time of infection. The compound C910 inhibited SARS-CoV-2 infection with an IC_50_ of 0.74 ± 0.07 µM (Figure 6B). Collectively, these results widen the protective spectrum of C910 against two RNA viruses that proceed through the endolysosomal pathway.

Both C910 and the PIKfyve inhibitor Apilimod induced swelling of EEA1-positive early endosomes (Sup. Figure 6A) (53). This suggested a possible PIKfyve inhibition in C910 action. Nevertheless, we found that Apilimod had no protective effect on CNF1-mediated degradation of Rac1 (Sup. Figure 6B) and conversely C910 did not inhibit PIKfyve kinase activity in vitro (Sup. Figure 6C). In conclusion, C910 bears broad anti-infective properties through a molecular mechanism different from that of Apilimod.

### C910 accumulates in lung tissues and protects mice against lethal Influenza

The pharmacokinetic (PK) parameters and pulmonary biodistribution of C910 following intraperitoneal administration was first defined. Supplementary Figure 7A shows that with a dose of 10 mg/kg, the plasmatic concentration of C910 reached a maximum (C_max_) of 164 ± 33 nM at 15 min, followed by a sharp drop to a prolonged plateau of about 25 nM for at least 24 h. C910 displays a half-life (T_1/2_) of 28.5 ± 4.3 h in the plasma, indicating a long exposure and slow elimination from the organism. Following a single intraperitoneal injection of C910 at 20 mg/kg/mouse, we estimated its concentration in lung tissues to 100-116 µM at 1 h and 61-110 µM at 4 h. We pursued by investigating the effect of intraperitoneal injection of 20 mg/kg of C910 on major blood chemistry parameters, e.g. markers of hepatic, pancreatic, renal and cardio-muscular functions using an approved field portable clinical device. No toxicity and no statistically significant differences were found between treated and vehicle control animals 24 h after administration (Sup. Figure 7B). Thus, despite low plasmatic concentrations, C910 accumulates in lung tissues with no recorded toxic effects and reaches for several hours effective inhibitory concentrations against IAV H1N1.

**Figure 7.**
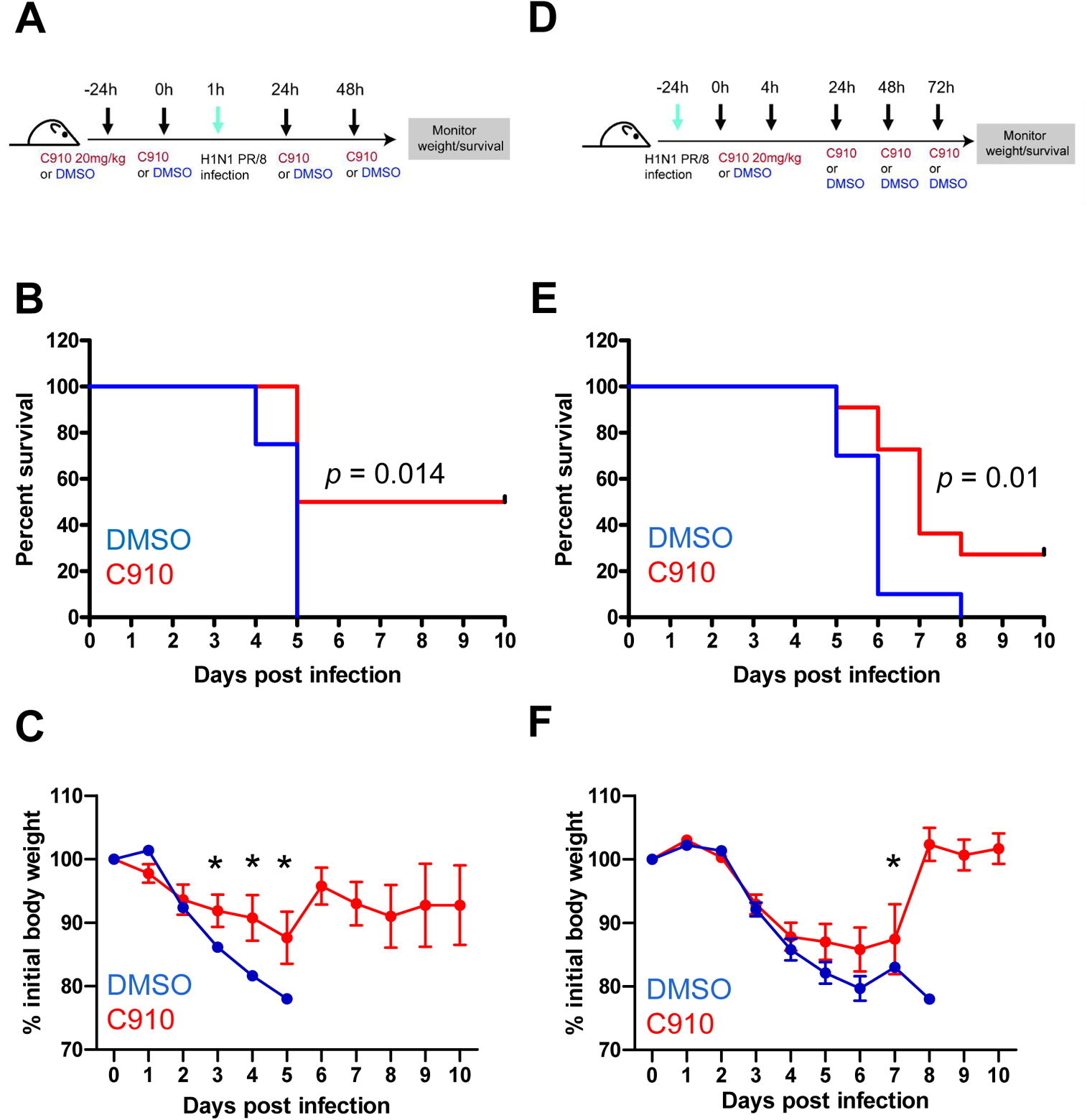
C910 protects mice against IAV H1N1 infection. A and D: time-line representation of the procedures. B-C: Survival curve for mortality (B) or weight loss (C) following intranasal infection with PR/8, as described in (A) are shown for mice treated with C910 (*n* = 8) or vehicle control DMSO (*n* = 8). E-F: Survival curve for mortality (E) or weight loss (F) following intranasal infection with PR/8 as described in (D) are shown for mice treated with C910 (*n* = 11) or vehicle control DMSO (*n* = 10). Mantel-Cox test was performed to compare the survival curve between groups injected with DMSO and C910. Weight loss data are shown as mean ± s.e.m, * *p* < 0.05, multiple *t*-test.

We then proceeded to proof of concept experiments starting with a prophylactic setting. C57BL/6 mice were intraperitoneally treated with C910 and then challenged with an infectious IAV (H1N1 PR/8 strain) (Figure 7A). At a low dose of virus known to induce weight loss and subsequent mortality, we recorded a complete protection of C910-treated mice, while the vehicle control-treated animals had to be euthanized during the course of infection due to excessive weight loss or serious symptoms (Sup. Figure 7C and 7D). When animals were challenged with a higher dose of virus in the same experimental setting, we recorded that C910 treatment improved mice survival (Figure 7B and 7C). Indeed, half of C910 treated-mice survived while all of the vehicle-treated mice had to be euthanized, starting at day 5. In a therapeutic setting (Figure 7D), repeated injections of C910 starting 24 hours after viral infection significantly increased mice survival and attenuated weight loss (Figure 7E and 7F). Taken collectively, these findings document a beneficial effect of C910 on mice infected with the influenza A virus, which can be attributed to the accumulation of C910 in lung tissues at an effective dose defined *in vitro*.

We conclude that C910 displays an extended protective spectrum against eight bacterial AB toxins and two viruses via an original molecular mechanism of action on early endosome sorting functions.

## DISCUSSION

We have isolated by HTS a piperidinamine-derived chemical compound that protects host cells against a wide spectrum of structurally unrelated AB toxins hijacking endolysosomal or retrograde membrane trafficking pathways. Mechanistically, we establish that C910 affects the morphology of Rab5/EEA1-positive endosomes and the routing of CNF1 and STxB to late endosomes and to the trans-Golgi network (TGN), respectively. We leveraged C910 properties to protect cells against infection by UPEC, IAV and SARS-CoV-2, and to confer protection of mice against IAV subtype H1N1 as well.

We show that direct monitoring of Rac1 cellular levels in response to CNF1 intoxication coupled to orthogonal screens allow the identification of a chemical agent acting on the two major endosomal pathways hijacked by AB toxins and several viruses. Direct measurement of the protective effect of chemical compounds on the target of toxins likely avoid identification of inhibitors acting indirectly via rescuing the downstream cytotoxic effects of specific toxin’s catalytic action. For example, the autophagy inhibitor 3-MA reverted autophagy-mediated cell death triggered by Stx2 via ER-stress pathway (54). Interestingly, we show that including orthogonal screens with DT and Stx1 is a stringent procedure that nonetheless allows the identification of a compound targeting the two major vesicular pathways hijacked by AB-like toxins, as compared to other anti-toxin chemicals identified to date (Sup. Table 2). These compounds comprise EGA and ABMA that were selected from HTS against the lethal toxin from *Bacillus anthracis* and the plant toxin ricin, respectively (2, 4). Retro-2 was selected according to the same procedure than ABMA, showing protection against ricin and Stx toxins (22). As summarized in the supplementary Table 2, these inhibitors display a specificity of action on a large panel of toxins trafficking through one of the two major pathways, as opposed to the broader spectrum of action of our newly discovered C910 compound. For example, C910 is able to inhibit Stx1/2, which are excluded from the anti-toxin spectra of ABMA and EGA. Moreover, Retro-2 is devoid of a protective effect against endolysosomal based-toxins, e.g. DT and CNF1 (unpublished data).

We attribute a vesicular traffic disruptor function to C910 on EEA1-positive early endosomes. Indeed, C910 does not affect the binding and endocytosis of CNF1 and Stx1 while it induces the swelling of Rab5/EEA1-positive early endosomes and alters their sorting functions. C910 affects the morphology of early endosome, in this window of treatment without inducing detectable changes on the subcellular distribution and morphology of other organelles studied. Diphtheria toxin translocation occurs primarily at the level of matured early endosomes (27). Our findings suggesting that C910 corrupts early endosome maturation are consistent with the recorded protection against DT that exits from matured early endosomes. Alteration of early endosome morphology by C910 resembles the phenotype induced by inhibition of the phosphoinositide 5-kinase PIKfyve that synthesizes phosphatidylinositol 3,5-bisphosphate (PtdIns(3,5)P_2_) from PtdIns(3)P (55). Of note, the knock-down of PIKfyve also affects the retrograde trafficking from early endosome to the TGN, and PIKfyve inhibitor shows cell protection against SARS-CoV-2 (51, 56, 57). Nevertheless, PIKfyve inhibition by Apilimod did not protect cells against CNF1, while it triggered early endosome enlargement at nanomolar range. Moreover, C910 has no inhibitory effect on recombinant PIKfyve kinase activity *in vitro*. Together, these data indicate that C910 displays an original molecular mechanism of action, different from that of Apilimod, although both molecules affect EEA1-positive early endosomes and display anti-infective properties. Characterization of the host target of C910 will thus contribute to define key steps in the vesicular trafficking exploited by numerous bacterial toxins and viruses.

We have previously screened 1,120 off-patent drugs from the Prestwick Chemical Library for their capacity to protect cells against CNF1-induced Rac1 depletion for drug repurposing. Nevertheless, the identified compounds did not allow further development in the clinic. Indeed, their antitoxin activities required concentrations higher than their safety range, and in some instances, they had effects on multiple cellular organelles (58, 59). The extended spectrum of protection conferred by C910 here points to its valuable pharmacological properties as much as it displays animal protection against IAV H1N1 with no recorded toxicity. As found for UPEC, C910 might protect cells from the pathogenicity triggered by bacterial pathogens responsible for severe or highly prevalent infections with limited susceptibility to antibiotics. This also relates to recurrent infections caused by *Clostridium difficile*. Development of host-targeted therapy is of particular interest against infections caused by Shiga toxin (Stx)-producing *E. coli* (STEC) as adverse effects cannot be fought by antibiotics (60) or CNF1-producing UPEC for which anti-microbial resistance is a growing concern (5). Interestingly, we show that C910 displays a higher protection against Stx2 compared with Stx1. Each toxin hijack different early endosome-Golgi transport pathways but Stx2 is responsible for triggering the hemolytic and uremic syndrome as a consequence of STEC infection in humans (8). Owing to the synthesis of LCGTs, *Clostridium difficile* induces antibiotic and healthcare-associated diarrhea with roughly 10% mortality rate and a high recurrence rate of approximately 20%. There is an urgent need in identifying novel treatments targeting LCGTs pointing to the importance of further characterizing C910 mechanism of action (61). The broad-spectrum of protection conferred by C910 also endows this molecule with promise against toxins from *Bacillus anthracis*, a pathogen rarely responsible for human infections and which is not the object of large vaccination campaigns but of concern of misuse. Therefore, host-targeted therapy targeting host vesicular trafficking represents an alternative strategy in the event of diversion of such a deadly pathogen.

Previous studies have shown that host-targeted prophylactic strategy endows the anti-toxin inhibitors an expanded efficacy against viruses (2, 4, 62, 63). Nevertheless, this is the first time to our knowledge that an anti-toxin inhibitor exhibits prophylactic potential against influenza A virus H1N1. In pilot experiments, we recorded a protective response induced by C910 in the group of male mice contrary to females (data not shown). This bias likely relates to the reported exacerbation of IAV-associated pathogenesis in female mice (64). Moreover, differences of C910 efficacy between male and female mice may reflect differences of pharmacokinetics.

Owning to its capacity of targeting two major vesicular trafficking pathways exploited by bacterial AB toxins and several viruses, C910 is a valuable tool to study membrane trafficking and help in the development of anti-infective compounds. The rapid evolution of the influenza virus complicates effective vaccine design therefore small molecule inhibitors for the treatment of viral infections are urgently needed. There is a number of antivirals in the clinic or in development, many of them targeting viral proteins. However, resistance against these antivirals rapidly emerges due to mutations in viral proteins (51, 65). Thus, the principle of new host-directed antitoxin and antiviral molecules such as C910 deserves further studies.

## MATERIALS AND METHODS

### Cell culture and bacterial toxins

Human umbilical vein endothelial cells (HUVECs) (PromoCell, Heidelberg, Germany) were cultured as described (24). HeLa, L929, Vero, A549, Vero E6 and U2OS-ACE2 cell lines were cultured at 37°C in 5% CO_2_ in DMEM/GlutaMax (Invitrogen) supplemented with 10% heat-inactivated FBS (F9665, Sigma-Aldrich) and 1% penicillin/streptomycin (Invitrogen). DT(D0564) was purchased from Sigma-Aldrich, Stx1 (#161) and Stx2 (#162) from List Biological, CNF1 was purified as described previously (58); TcdA and TcdB were purified from *C. difficile* VPI10463 as described (66); protective antigen and lethal factor from *Bacillus anthracis* were purified as described (67); PE toxin was kindly provided by Bruno Beaumelle (UMR5236 CNRS, University of Montpellier, France); STxB was purified and labeled with Alexa Fluor 488, as described (68, 69).

### Antibodies and reagents

C910 was purchased from Chembridge (ID: 5454910, San Diego, CA, USA) or Synthenova (ID: SN0218L3, Hérouville Saint-Clair, France). The following products were purchased from the indicated commercial sources: L-[^14^C(U)]-leucine was from Perkin-Elmer; DMSO (D4540), Hoechst 33342 (B2261), Gelatin (G9361), Saponin (S-7900), Bafilomycin A1 (B1793), mouse anti-actin (ac-74), HIS-tagged B subunit of Shiga toxin 1(SML0655) and Apilimod (A149227) from Sigma; rabbit anti-EEA1 (#3288), rabbit anti-Rab7 (#9367), rabbit-anti-his tag (#12698) from Cell Signaling; rabbit anti-Rab7 (ab137029), mouse anti-Giantin (ab37266) and Cathepsin B Activity Assay Kit (ab65300) from Abcam; mouse-anti-6x-his (R930-25 [clone 3D5]), DQ^TM^ Red BSA (D-112051) from ThermoFisher Scientific; mouse-anti-EEA1 (BD610457), mouse-anti-Rab5 (610724), mouse anti-Rac1 (610651 [clone 102]), mouse anti-Rab4 (610888), mouse anti-Lamp1 (611043) was from BD Bioscience; mouse anti-TGN38 (sc-101273), rabbit anti-MEK2 (SC-524) from Santa Cruz; rabbit anti-Calnexin (ADI-SPA-860-D) from Enzo; mouse-anti-LBPA (Z-PLBPA) was from Echelon Biosciences; Paraformaldehyde (PFA) (15710) from Electron Microscopy science. Fluorescence-tagged secondary antibodies were purchased from Thermofisher and used at 1:500 dilutions: Alexa Fluor 594 donkey anti-mouse (A-21203), Alexa Fluor 594 donkey anti-rabbit (A-21207), Alexa Fluor 488 donkey anti-mouse (A-21202), Alexa Fluor 488 donkey anti-rabbit (A-20206), Alexa Fluor 647 donkey anti-mouse (A-31571). Horseradish peroxidase (HRP)-conjugated goat anti-mouse (P0399) or swine anti-rabbit secondary antibodies (P0447) were from Dako.

### Orthogonal screening pipeline

All steps of compound screening were performed in gelatin-coated Nunc MicroWell 96 Well opaque plates (Thermo Scientific) seeded with 20,000 HUVECs per well. During the first round of screen, we tested 16,840 compounds from the ChemBridge DIVERSet library at a working concentration of 50 μM in the presence of CNF1 10 nM during 6 h (Z’ values > 0.5).

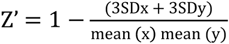

Rac1 cellular levels were determined by direct immunolabeling of cells, as followed. Fixation of HUVECs was performed in 4% PFA, 15 min. Permeabilization and saturation were performed concomitantly in D-PBS supplemented with 1% donkey serum, 0.1% Triton X-100 and 0.05% Tween-20 during 1 h. Cells were next labelled with an anti-Rac1 mouse antibody [clone 102] for 1 h and revealed with Alexa Fluor 488-coupled anti-mouse secondary antibody. Rac1 immunostaining was quantified using a Cytation 5 reader (BioTek) in a fluorescence intensity mode (λ_Ex_= 485 nm; λ_Em_= 520 nm) with a 3 x 3 area scan mode per well. Hit compounds were cherry-picked and reevaluated following the same procedure with another secondary antibody coupled with Alexa Fluor 594 (λ_Ex_= 590 nm; λ_Em_= 620 nm) to exclude auto fluorescent molecules. Selected hits were then reordered and freshly prepared for a third round of screen. Defined hit compounds were further filtered from pan-assay interference compounds (PAINS), i.e. unstable molecules, irreversible modifiers or compounds that are frequently active in other screens (25). Eleven compounds were shortlisted as robust inhibitors of CNF1-mediated degradation of Rac1 and went through a series of two orthogonal screens to define their protective effects first against DT and then Stx1. Compound N-(3,3-diphenylpropyl)-1-propyl-4-piperidinamine, here referred as C910, passed all the screens. Of note, C910 and the unrelated (S)-2-amino-N-(1-((5-chloro-1*H*-indol-3-yl)methyl)piperidin-4-yl)-3-(1*H*-indol-3-yl) propanamide (referred to as API or SN0209) have been isolated for their capacity to alter the interaction of SOS with p21-Ras GTPase *in vitro* (70). Co-treatment of cells with API ranging from 5 to 10 μM did not protect cells against CNF1 ruling out a possible protective action of C910 through the SOS/Ras signaling (not shown).

### Evaluating efficacy of C910 against intoxication

Protein synthesis inhibition was measured as described (22). Briefly, cells were plated overnight in 96-well Cytostar-T microplates with scintillator incorporated into the polystyrene. Cells were then challenged with increasing doses of toxins for the indicated periods of time (DT: 6 h; Stx1, Stx2, PE: 18 h) in presence of vehicle or various concentrations of C910. The medium was replaced with DMEM supplemented with L-[^14^C(U)]-leucine at 0.5 μCi/ml for an additional 3 h (Stx1, Stx2, PE) or 15 h (DT). Protein synthesis was determined using a Wallac 1450 MicroBeta scintillation counter (PerkinElmer). All values are expressed as means of duplicates ± s.e.m., data were fitted using the nonlinear regression dose-response and the goodness-of-fit for toxin alone (carrier) or with drugs was assessed by *r^2^* values and confidence intervals with Prism ver. 8 software (GraphPad, San Diego, CA).

Cytotoxic effects of the LCGTs TcdA and TcdB on Vero cells (Sigma-Aldrich) were evaluated by measure of cell rounding. Vero cells seeded in 96-well plates at a density of 10,000 cells per well one day before, were intoxicated with 8 cytotoxic units (CU) of TcdA or TcdB for 8 h in the absence or presence of C910 at 0, 5, 10 and 20 µM. Cell rounding was visualized by phase-contrast microscopy. In parallel, cell lysates from intoxicated cells were examined for glucosylated-Rac1 that no longer react to anti-Rac1 monoclonal antibody [clone 102].

The activity of LT from *B. anthracis* was assessed by determination of MEK2 cleavage. Cell lysates from HUVECs intoxicated with two concentration of LT (0.3 μg/ml protective antigen (PA) + 0.1 μg/ml Lethal factor (LF) or 0.03 µg/ml PA + 0.1 µg/ml LF) in the absence or presence of C910 at 40 µM for 1, 2, 4, 8 h were collected and examined for MEK2 protein levels by immunoblot. GAPDH was used as the loading control.

### Western blot analysis

Samples were size-separated via SDS-PAGE, transferred to Immobilon-P PVDF membranes (Millipore) and labelled by primary antibodies. Signals were revealed using horseradish peroxidase-conjugated goat anti-mouse or swine anti-rabbit secondary antibodies (DAKO) followed by chemiluminescence using Immobilon® Western (Millipore). The chemiluminescent signals were recorded on a Syngene PXi imager (OZYME), and the data were quantified using Fiji software (NIH).

### Binding of CNF1 to cells analyzed by FACS

Recombinant CNF1 toxin was fluorescently labeled with Cy3 (Cy®3 Mono 5-pack, GE healthcare, PA23001) according to the vendor’s recommendations. Briefly, purified toxin preparations (1 mg/ml) were incubated with the Cy3 dye in 100 mM bicarbonate buffer for 18 h at 4°C and applied to dialysis to separate the toxin from the free dye. CNF1-Cy3 (16 nM) was absorbed to the cells at 4°C for 30 min and cells processed for FACS analysis using a BD LSRFortessa™ flow cytometer. Data were analyzed with FlowJo (BD Bioscience).

### Fluid-phase transport to lysosomes

HUVECs were seeded in fibronectin pretreated-µ-Slide 8 Well Glass Bottom (80827, ibidi) one day before. Cells were pulsed with DQ-BSA (DQ^TM^ Red BSA) at a concentration of 10 μg/ml for 30 min (37°C, 5% CO_2_). The cells were then washed three times with PBS before being treated with compounds in the complete medium at 37°C for the indicated time, nuclei were labeled with Hoechst 33342. Fluorescence of live cells were acquired with a Perkin Elmer Ultraview spinning disk confocal microscope using 60X/1.2 NA water objective.

### Immunocytochemistry, confocal imaging and image quantification

Immunocytochemistry was performed on cells fixed with 4% PFA in PBS for 15 min at room temperature. Next, cells were permeabilized with 0.1% Saponin in PBS for 10 min and blocked with 0.2% gelatin in PBS. Immunolabeling with indicated antibodies was performed in 0.05% Saponin and 0.1% gelatin in PBS Buffer. Images were acquired using a Perkin Elmer Ultraview spinning disk confocal microscope using 60X/1.2 NA oil objective. Fiji ImageJ software (National Institutes of Health) was used for image processing and quantification.

Measures of co-localization between two fluorescence signals of interest are expressed as Pearson’s coefficient correlation or Mander’s overlap coefficient. Cells in each condition were randomly selected and analyzed under the same threshold. Briefly, each individual cell was selected from a single mid-z stack of the confocal image by hand-drawing a “region of interest” (ROI) from Fiji software. The two channels of fluorescence were next separated and analyzed with JACop plug-in.

Measures of the size of EEA1-positive vesicles were performed from the single mid-z stack of confocal images that were next quantified with the Analyze particles tool from Fiji Software. Threshold values were maintained throughout each experimental set.

### Bacterial infection experiments

The UTI89 uropathogenic *E. coli* (UPEC) mutant strain was described previously (71). *Salmonella enterica* serovar Thyphimurium strain SL1344 was kindly provided by Jost Enninga (Institut Pasteur, France). Bacteria internalization into HUVECs was assessed using the gentamycin protection assay. Briefly, HUVECs were seeded on 12-well plates at the density of 250.000 cells/well overnight (triplicate wells). When indicated, 1 nM CNF1 toxin was added after 30 min pre-treatment with 15 μM C910. Exponentially growing UPEC (OD_600_ = 0.6 in LB) or *Salmonella* (OD_600_ = 0.6 in LB with 0.3 M NaCl) were added onto cells at MOI 100, followed by 20 min centrifugation at 1,000 x g. Cell infection was performed for 10 min at 37°C, 5% CO_2_. Infected cells were washed three times with PBS and either lysed for cell-bound bacteria measurements or incubated another 30 min in the presence of 50 μg/ml gentamicin before lysis for internalized bacteria measurements. Cells were lysed in PBS + 0.1% Triton X-100 and serial-diluted bacteria plated on LB-agar supplemented with 200 μg/ml streptomycin or ML-agar for CFU counting.

### IAV infection

For single-cycle IAV infection assays, A549 cells (70,000/ well) were infected with a reporter H1N1 A/WSN/33 (H1N1_WSN_ PB2-2A-mCitrine), at a MOI of 5 PFU/cell for 6 h. Cells were fixed in 4% PFA and analyzed by flow cytometry (Attune NxT; Thermo Fisher Scientific) as described in (50). For multicycle growth assays, A549 cells were infected with a seasonal A/Bretagne/7608/2009 (H1N1_pdm09_) strain adapted to human cell lines (50), at the MOI of 10^−3^ infective particles/cell and the production of infectious particles in the culture supernatant was determined at 24 h using a standard plaque assay on the highly sensitive canine MDCK-SIAT cells, as described previously(72).

### SARS-CoV-2 infection

U2OS-ACE2 GFP1-10 and GFP 11 cells, also termed S-Fuse cells, become GFP+ when they are productively infected by SARS-CoV-2 (49). Cells were mixed (ratio 1:1) and plated at 4×10^3^ per well in a μClear 96-well plate (Greiner Bio-One). The following day, cells were incubated with the indicated concentrations of C910 for 2 h at 37°C. SARS-CoV-2 (BetaCoV/France/IDF0372/2020) (MOI 0.1) was then added. 20 h later, cells were fixed with 2% PFA, washed and stained with Hoechst 33342 (dilution 1: 1,000, Invitrogen). Images were acquired with an Opera Phenix high content confocal microscope (PerkinElmer). The GFP area and the number of nuclei were quantified using the Harmony software (PerkinElmer). The percentage of inhibition was calculated using the GFP area value with the following formula: 100 x (1 – (value with IgA/IgG – value in “non-infected”)/(value in “no IgA/IgG” – value in “non-infected”)).

### *In vivo* IAV infection model

All mice were housed under specific-pathogen-free conditions at Seattle Children’s Research Institute and all animal experiments performed at Seattle Children’s Research Institute were approved by the Institutional Animal Care and Use Committee (IACUC00580). A pilot experiment showed a protective effect of C910 in male animals therefore we used male mice for further experiments (data not shown). Viral challenges were carried out on groups of 9~10 week-old C57BL/6 male mice (Charles River Laboratories). Animals were treated by intraperitoneal injection with either 20 mg/kg of C910 or vehicle control DMSO as described in the figures. Anesthetized mice were infected intranasally with the IAV H1N1 PR/8 strain (Charles River Laboratories) diluted in 25 μL PBS, at a high dose (1.5×10^5^ PFU/mouse, Figure 7A-C) and at a low dose (5×10^3^ PFU/mouse, Sup. Figure 7D-E) in prophylactic setting as well as a dose of 9.5×10^3^ PFU/mouse (Figure 7D-F) in treatment setting. Mice were monitored daily for weight loss and any other signs of disease.

### Statistics

All data are presented as mean ± s.d. or mean ± s.e.m., unless specified in the figure legends. Student’s *t-*tests were applied to determine the statistical significance between two datasets and one-way ANOVA was used to perform analyses between more than two datasets, unless specified in the figure legends. A value of *p* ≤ 0.05 was considered significant (denoted * *p* ≤ 0.05, ** *p* < 0.01, *** *p* < 0.001, **** *p* < 0.0001). Graph Pad Prism Software 8 was used to produce the graphs and to perform statistical analyses.

## ACKNOWLEDGEMENT

We thank Cezarela Hoxha, Laurent Audry, Yuen-Yan Chang, Sylvain Meunier (Institut Pasteur, Paris, France) and Imène Belhaouane (Institut Pasteur, Lille) for technical involvement as well as Caroline Stefani (Benaroya Research Institute, USA), Daniel Ladant (Inst. Pasteur) and Jean-Nicolas Tournier (IRBA) for fruitful discussions. This work was funded by the joint ministerial program of Research and Development against Chemical, Biological, Radiological, Nuclear and Explosive risks (CBRNE), a grant from the Agence Nationale de la Recherche (ANR-19-ASTR-0001), Labex IBEID (ANR-10-LABX-62-IBEID), the French Alternative Energies and Atomic Energy Commission (CEA). N.M., L.C., F.V. and E.P. were supported by DGA-MRIS/AID scholarships with CEA and Pasteur, respectively. SIMoS and SCBM are members of the Laboratory of Excellence LERMIT, supported by a grant from the Agence Nationale de la Recherche (ANR-10-LABX-33).

## COMPETING INTERESTS

All authors do not have any financial or other interests related to the submitted work.

## AUTHOR CONTRIBUTIONS

E. L. and D.G. coordinated the research; J.B. directed HTS; N.M., J-C.C. and J.B. operated the HTS and analyzed results; Y.W. and E.P. performed cell biology experiment; L.S., Y.W., N.M., E.P., and V.K. performed measurements of C910 efficacy on toxins; IAV H1N1 experiments were performed by C.D and S.V. (*in vitro*) and S.G and M.A (*in vivo*); SARS-CoV-2 experiments was performed by F.G. and O.S.; S.P. and A. Mettouchi. conceived and operated bacterial invasion experiments; M.S. and Y.W. performed electron microscopy experiments; L.C., P.C., R.T., P. Barbe., M.K., P. Bordin., A. Machelart., V.S. and F.T. contributed to pharmacokinetics and biodistribution studies, M.K. and R.T. established C910 concentration in biological samples; G.C. and B.P. performed *in vitro* PIKfyve activity measurements; M-R.P. and L.J. supplied LCGTs and STxB respectively; J-C.C. and F.V. supplied chemical compounds; Y.W., D.G. and E.L. drafted the manuscript. All authors read and approved the manuscript.

## Materials and Methods

### Determination of EC_50_s of C910 for DT, Stx1, Stx2 and PE

The mean percentage of protein biosynthesis was determined and normalized from duplicate wells. All values are expressed as means ± s.e.m. Data were fitted with Prism v8 software (Graphpad Inc., San Diego, CA) to obtain the toxin concentrations giving 50% protein synthesis inhibition in absence or presence of compound (IC_50 drug_ vs IC_50 DMSO_). The protection factors R (R = IC_50 drug_/IC_50 DMSO_) were determined by the software’s nonlinear regression “dose-response EC_50_ shift equation”. For each concentration of C910, a percentage of protection was determined from R values (% protection = (R-1) / (Rmax − 1) × 100, with Rmax corresponding to the highest value of R in the series). The concentration of C910 was plotted against the corresponding percentage of protection of cells and the 50% efficacy concentration (EC_50_) was calculated by non-linear regression.

### Determination of CC_50_s of C910 on HUVECs and HeLa cells

Cells seeded in Corning™ 96 well Flat Clear Bottom Black Microplates (#3603) were incubated with increasing doses of C910 for 6 h, medium was replaced with fresh medium including 10% of Alamar Blue (final concentration 1:10). Then the fluorescence (540/590 nm) was measured 1h later by Cytation 5 cell imaging multi-mode reader (BioTek). The mean percentage of toxicity was determined from duplicate wells. All values are expressed as means ± s.e.m from one representative experiment. Data were fitted with Prism v8 software (Graphpad Inc., San Diego, CA) to obtain the concentrations giving 50% toxicity.

### Electron microscopy

After the uptake of BSA-gold (Cell Microscopy Core, Universitair Medisch Centrum Utrecht, Netherland), HUVECs were fixed at room temperature for 2 h in 2.5% glutaraldehyde in PHEM buffer, pH 7.2 (60 mM 1,4 piperazine diethylsulfonic acid (PIPES), 25 mM N-2-hydroxyethylpiperazine Nl-2-ethane sulfonic acid (HEPES), 10 mM EGTA, 2 mM MgCl2, pH 7.2). After washes with PHEM, cells were post-fixed with 1% osmium and 1.5% potassium ferricyanide (Merck, Darmstadt, Germany) in PHEM before dehydration in a graded series of ethanol and infiltration with epoxy resin. Resin was polymerized at 60°C for 48 h. Sections with a nominal thickness of 70 nm were obtained with a UC7 (Leica Microsystems, Vienna, Austria) and collected on formvar, carbon coated 200 mesh copper grids (Electron Microscopy Sciences, Hatfield, PA, USA). Sections were contrasted with 4% uranyl acetate and Reynold’s lead citrate and observed with a Tecnai spirit (Thermofisher, Eindhoven, Netherlands) operated at 120 kV. Images were acquired using a Gatan Ultra Scan™ 4000 digital camera (Gatan, Pleasanton, USA).

### *In vitro* PIKfyve activity

Recombinant PIKfyve production and activity measurements were performed as described ^1^. Briefly, PIKfyve was incubated with DMSO or compounds in the presence of PtdIns3P/PE lipid vesicles for 15 min at room temperature, prior to addition of [ɤ^32^P]-ATP and incubation at 37°C for 30 min. After lipid extraction and thin layer chromatography, the [^32^P]-PtdIns(3, 5)P_2_ produced by PIKfyve was visualized by autoradiography.

### C910 dosage and quantification by LC-MS/MS for pharmacokinetics and biodistribution studies

Two stock solutions of C910 standard and another compound used as internal MS standard (IS) were prepared at concentrations of 10 mM in DMSO. For establishing calibration plots, known amounts of C910 were spiked into control mouse plasma or lung homogenates. To eliminate plasmatic proteins and other endogenous compounds, C910 was extracted from plasma samples by two extractions with acetone. This involves addition of 4 volumes of ice-cold acetone and incubation for 1 h at −20°C. After centrifugation at 15,000 x g for 10 min, the supernatants containing C910 were collected. For each sample, both acetone supernatants were combined and dried at 40 °C for 3 h. Dried extracts were finally re-suspended into 25% acetonitrile (ACN) in water containing 0.1% formic acid (FA) for analysis by LC-MS/MS Multiple Reaction Monitoring (MRM) mode.

Mouse lungs were homogenized using a Precellys/Cryolys Evolution homogenizer from Bertin, France, according to the manufacturer’s recommendations using the CK28 ceramic beads 2 mL lysing kit. One hundred mg of organ in 400 µL of cold PBS were grinded at 4 °C by 3 to 6 cycles of 30 sec at 5,000 rpm with 15 s pause between each cycle until obtaining a smooth homogenate (3 animals per time point, two independent experiments).

To extract C910 from lung homogenates, samples were washed twice with ice-cold PBS (vortex, centrifugation at 15,000 x g for 10 min, elimination of the supernatant). Then, the pellets were resuspended, vortexed and centrifuged twice in DMSO to extract C910 and both supernatants were combined. A fraction of this DMSO supernatant extract was diluted with 25% ACN in water containing 0.1% FA and analyzed by LC-MS/MS (MRM mode). The LC-MS/MS system consisted of an Agilent 1100 HPLC system on-line coupled to an Esquire-HCT ion Trap mass spectrometer (Bruker-Daltonics) equipped with an electrospray ionization (ESI) source. LC separation was carried out at a flow rate of 200 µL/min on a ThermoScientific reverse phase Accucore-150 C18 column (2.1 x 50 mm; 2.6 µm) with a linear gradient 0-100% B for 7 min with a mobile phase composed of 0.1% FA in water for A and 0.1% FA in ACN for B. ESI conditions and detection of C910 and products were optimized as well as for the internal MS standard IS spiked in samples for quantification. Fragmentation energy amplitudes were also optimized (SmartFrag off). MS detection was operated in MRM mode. The fragmentation transition followed for dosage of C910 was m/z 337.2 → 125.9. A second fragmentation transition was used to confirm the identity of C910 (m/z 337.2 → 238.0). The fragmentation transition followed for internal standard was m/z 436.0 → 284.9 (dosage) and m/z 436.0 → 254.9 (identity confirmation). Data Analysis software (Bruker Daltonics) was used for qualitative and quantitative processing of raw MS and MS/MS data. The ion chromatograms of MS/MS transitions of C910 as well as of the internal MS standard were extracted (EIC). These EIC peaks were integrated and the ratio C910/IS of the corresponding MS transitions enabled to plot calibration curves of C910 in plasma or lung homogenates and further quantity C910 in any of these biological matrices from mice treated with C910.

### *In vivo* pharmacokinetics (PK), biodistribution and plasmatic diagnostic

Animal care and surgical procedures were performed according to the Directive 2010/63/EU of the European Parliament, which had been approved by the Ministry of Agriculture, France. The project was submitted to the French Ethics Committee CEEA (Comité d’Ethique en Expérimentation Animale) and obtained the authorization APAFIS#10108-2017060209348158 v3. Animal protocols in CIIL (Center for Infection and Immunity of Lille) were approved by the Minister of Higher Education and Research after favorable opinion of the Ethics Committee (CEEA Nord-Pas de Calais n°75/APAFIS#10232-2017061411305485 v6). All experiments were performed in accordance with relevant named guidelines and regulations. Experiments were conducted on adult male C57BL/6J mice (23.5 ± 0.3 g) purchased from Janvier Labs (Le Genest St Isle, France). Mice were injected intraperitoneally with 10 mg/kg, i.e. 29,716 nmol/kg, of C910. Blood samples were collected at 5 min, 15 min, 30 min, 45 min, 1 hr, 6 h and 24 h after injection, on three distinct mice at each time. Quantification of C910 concentration contained in plasma at each time was performed by LC-MS/MS. PK parameters were calculated from these data with the non-compartmental analysis Linear up/Log down method, using the PKSolver add-in program. Plasmatic diagnostic was performed at 24 hrs after injection using the Piccolo® AmLyte 13 and the Piccolo Xpress® chemistry analyzer (Abaxis, USA). Levels of glucose, blood urea nitrogen, creatinine, total bilirubin, albumin, alanine aminotransferase, aspartate aminotransferase, creatine kinase, amylase, sodium, potassium, and calcium were determined in heparinized plasma of five mice injected with vehicle or C910. Quantitative data are shown as means, with error bars indicating the standard error of the mean (SEM).

## Figure legends

**Supplementary Figure S1.**
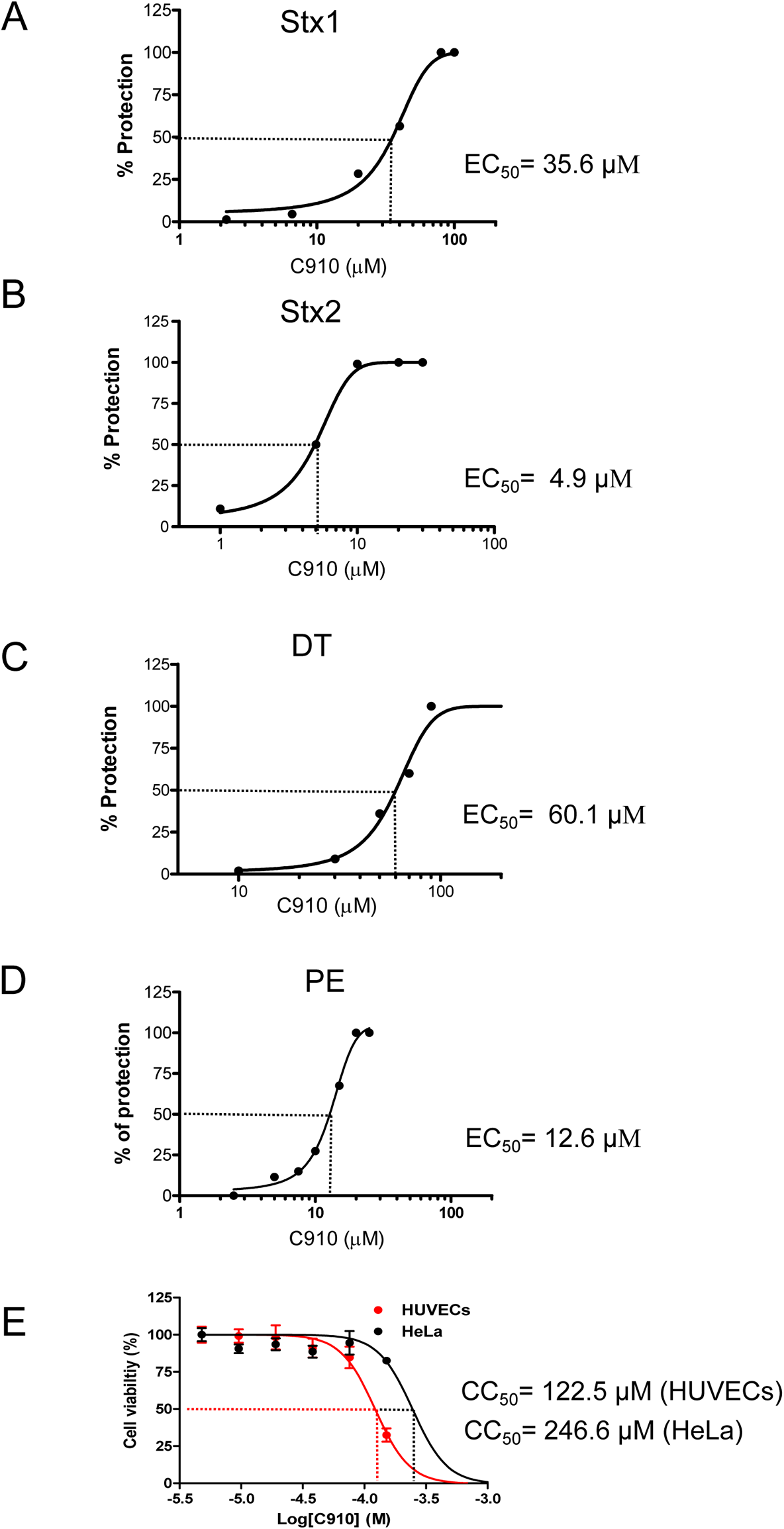
EC_50_s of C910 for DT, Stx1 and Stx2 and its CC_50_, corresponding to Fig 1C and E, F. Curves show different concentrations of C910 and the corresponding percentages of cell protection against Stx1(A), Stx2(B), DT(C) and PE(D). (E) Curves show different concentrations of C910 and the corresponding viabilities of HUVECs and HeLa from one representative experiment (n = 2).

**Supplementary Figure S2.**
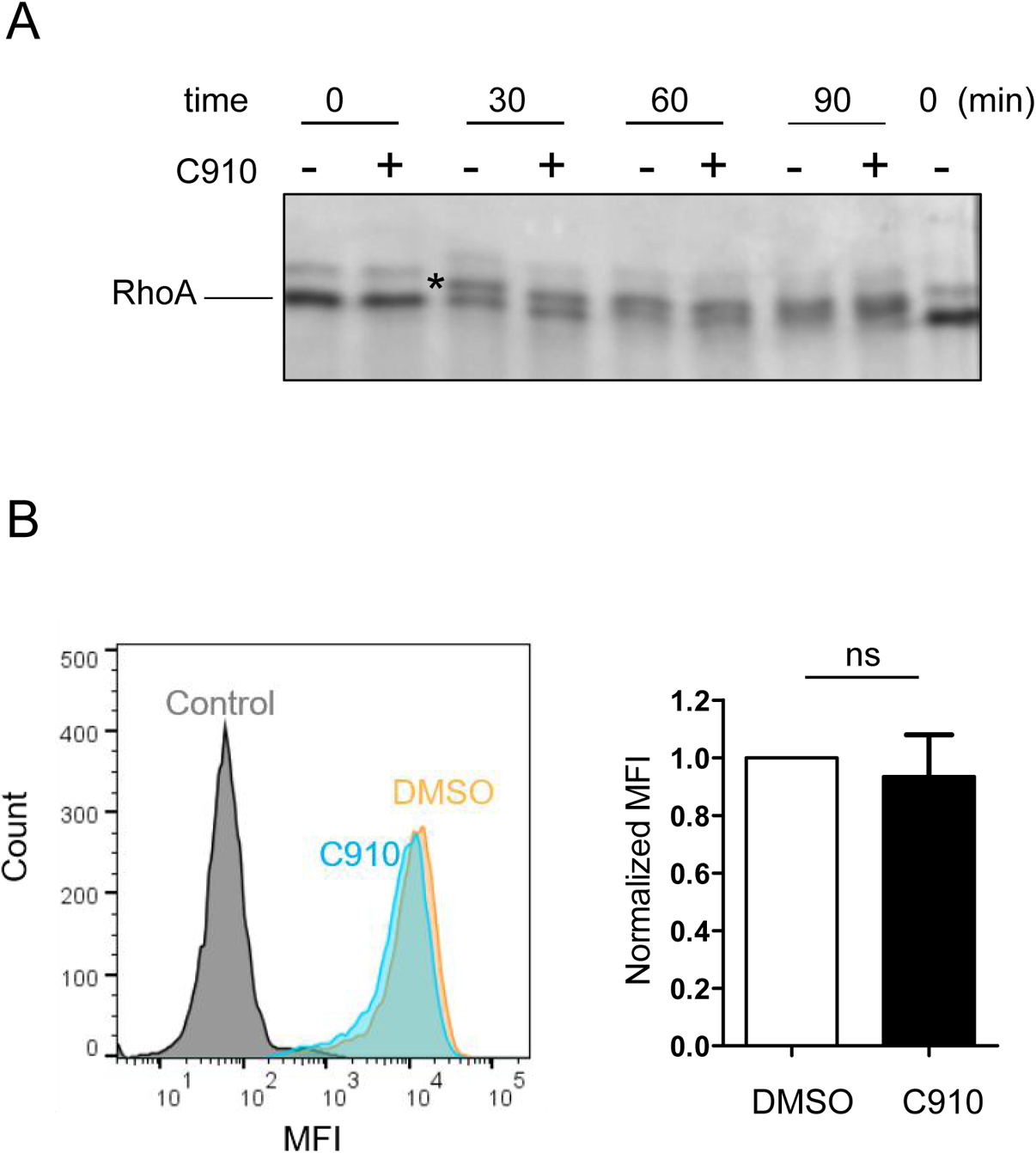
C910 does not affect CNF1 catalytic activity and STxB binding with membrane receptors. A: Recombinant RhoA protein (1 μg) was incubated with CNF1 (0.1μg) in the presence or absence of C910 (1 µM) for the indicated time. The time-dependent upper-shift of RhoA shows the absence of effect of C910 on CNF1 deamidase activity. B: HeLa cells were pre-treated with 40 µM C910 or DMSO for 1 h at 37°C. Next, 3 mM EDTA-dissociated cells were incubated with STxB-his on ice for 30 min in the presence of 40 µM C910 or DMSO vehicle and subsequently fixed with 4% PFA, labelled by anti-his antibodies and Alexa Fluor 488-conjugated secondary antibodies and then analyzed by Flow Cytometry. Signal of cells without incubation with STxB-his was used as background. MFI (Mean Fluorescence Intensity) of C910-treated cells was normalized to DMSO-treated cells values. The histogram represents mean ± s.e.m. of four independent experiments, ns: not significant, paired two-tailed *t*-test.

**Supplementary Figure S3.**
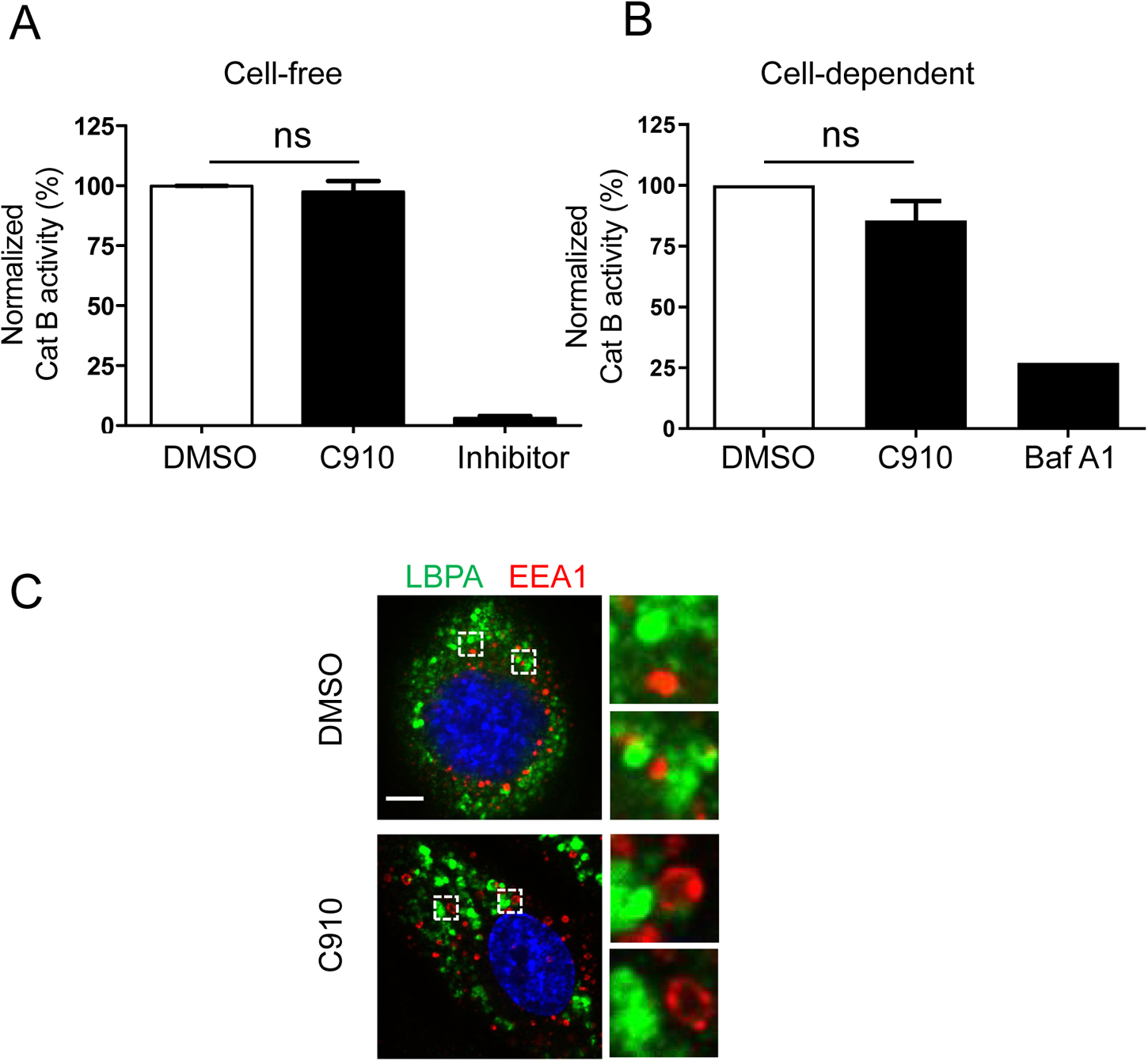
C910 does not inhibit cathepsin B activity. A: Cell-free assay. Cell lysates mixed with cathepsin B substrate labelled with fluorescent probe, and incubated 2 h with DMSO vehicle, 20 µM C910 or a control inhibitor, provided in the kit. B: Cell-dependent assay. Lysates from HUVECs treated with DMSO vehicle or 20 µM C910 for 2 h were examined. Cell lysate supplemented with Bafilomycin A1 (Baf A1) 200 nM was used as control of inhibition of acid-dependent cathepsin B. Fluorescent intensity (λ_Ex_/λ_Em_ = 400/505 nm) from cleaved fluorescent probe was determined on FluoStar Omega (BMG Labtech) and normalized to DMSO vehicle, as a control. Data represent the mean ± s.e.m. of three independent experiments. ns: not significant, paired two-tailed *t*-test.

**Supplementary Figure S4.**
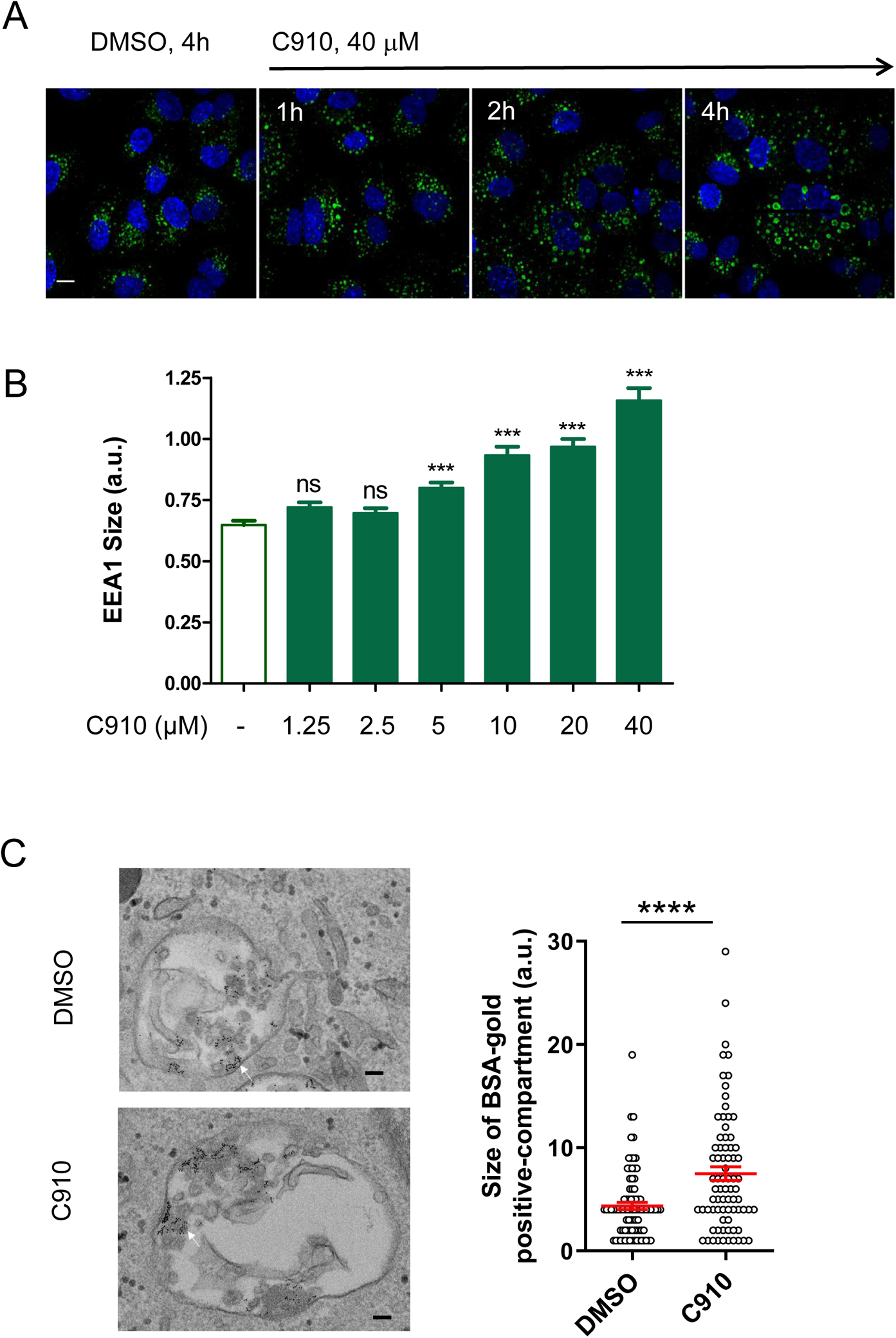
Dose and time-dependent enlargement of EEA1-positive compartments in C910 treated cells. A: HUVECs were incubated with C910 or DMSO for the indicated periods of time, fixed, permeabilized and labeled for EEA1. Representative images are shown from two replicate experiments. Scale bar, 10 μm. B: HUVECs were incubated with different doses of C910 or DMSO for 20 h and then processed as in (A). The size of EEA1-positive vesicles from at least 20 cells in each condition are quantified by Image J and shown as mean ± s.e.m. One-way ANOVA was performed overall (*p* < 0.0001) with Dunnett’s multiple comparison test to compare control and treated conditions, *** *p* < 0.001, ns: not significant. C: HUVECs were pre-treated with 20 µM C910 or DMSO vehicle for 2 h and then pulsed with BSA-gold for 15 min in the presence of C910 or DMSO vehicle. Cells were processed for electron microscopy. Representative electron micrographs are shown. White arrows indicate BSA-gold. Right, the size of BSA-gold positive compartments was quantified and data presented as mean ± s.e.m. for 87 (DMSO) and 77 (C910) compartments. Each point represents an individual compartment, **** *p* < 0.0001, unpaired two-tailed *t*-test. Scale bar, 100 nm.

**Supplementary Figure S5.**
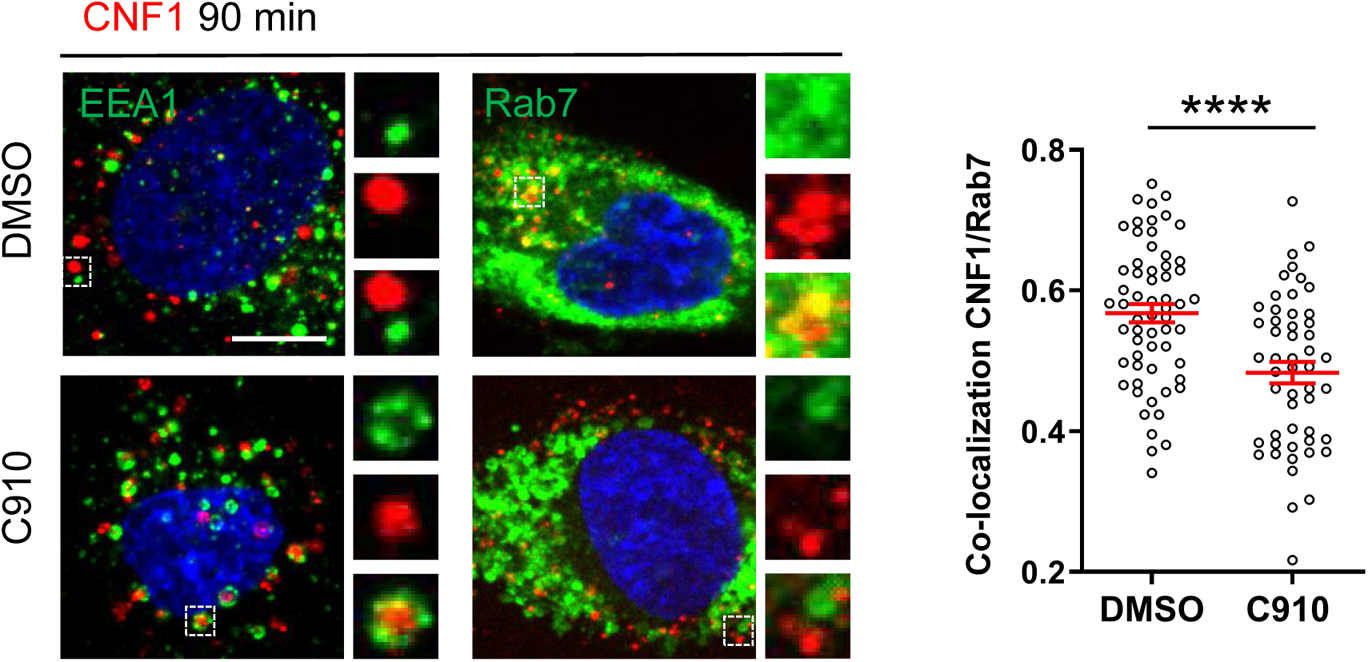
C910 affects the trafficking of CNF1, corresponding to Figure 5C. HUVECs were treated with 20 µM C910 or DMSO as Figure 5C. Right, Pearson’s coefficient of two fluorescence (CNF1-Cy3 and Rab7) was analyzed with Fiji software and JACop plug-in. Data are presented as mean ± s.e.m. for 60 (DMSO) and 51 (C910) cells from three independent experiments. Each point represents an individual cell, **** *p* < 0.0001, unpaired two-tailed *t*-tests. Scale bar, 10 µm.

**Supplementary Figure S6.**
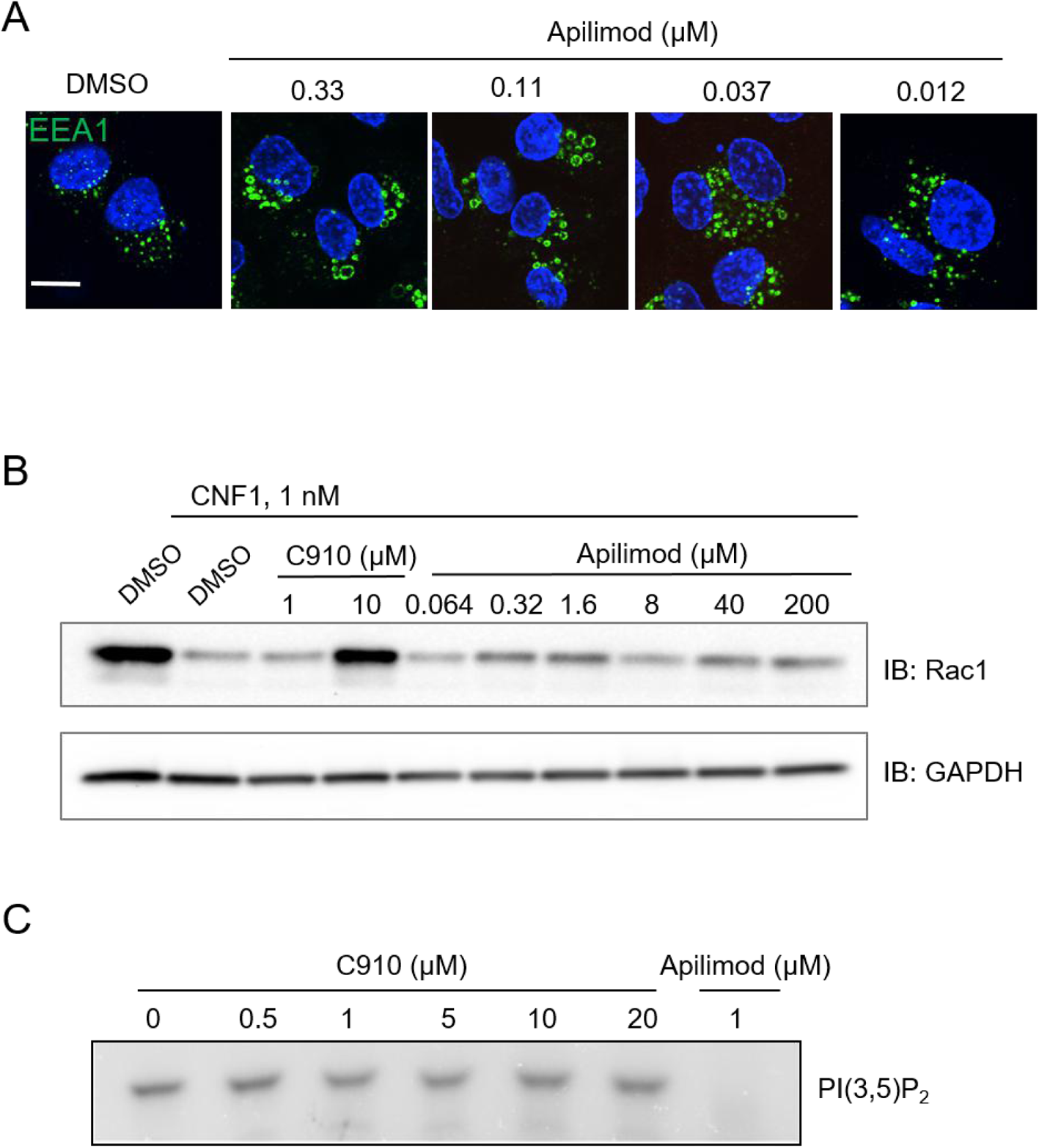
Distinctive mechanisms of action between Apilimod and C910. A: Representative images show EEA1 staining of HUVECs incubated with DMSO or Apilimod at the indicated concentration for 2 h. Scale bar, 10 μm. B: Immunoblots anti-Rac1 show levels of CNF1-mediated Rac1 degradation and protective effect of C910 treatment on Rac1 degradation. In contrast, Apilimod has no significant protective effect at a broad range of concentrations. One representative experiment, *n* = 2. HUVECs were pre-incubated for 30 min with indicated concentrations of compounds prior to addition of CNF for 6 h. Anti-GAPDH was used as loading control. C: Representative images show radioactive labelled PI_(3,5)_P_2_ produced by recombinant PIKfyve in the presence of DMSO or compounds. In contrast to Apilimod, increasing doses of C910 had no effect on recombinant PIKfyve lipid kinase activity. One representative experiment, *n* = 2.

**Supplementary Figure S7.**
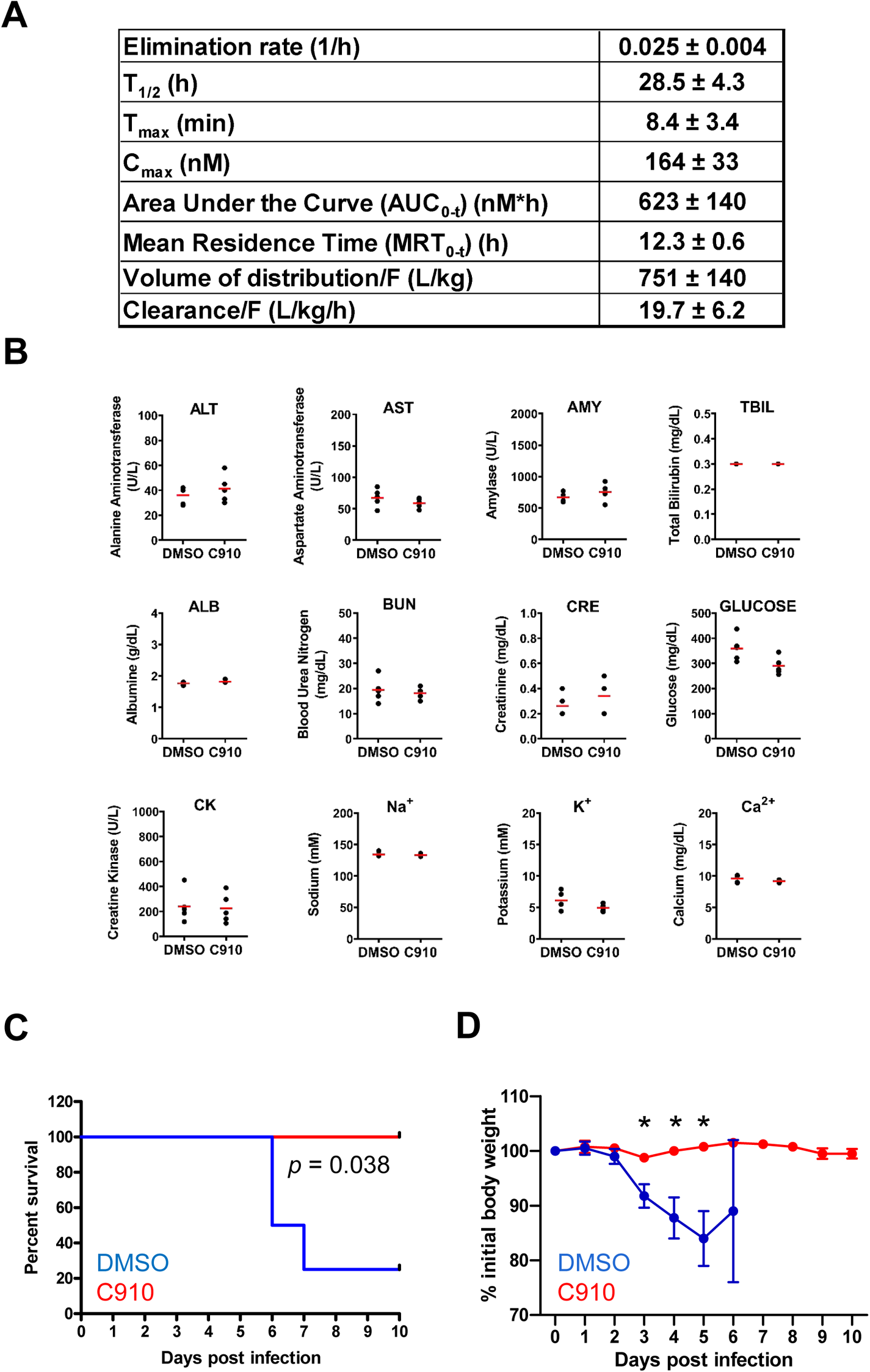
C910 PK, blood diagnostic and inhibition of IAV infection *in vivo*. A: PK parameters calculated for C910 following an intraperitoneal administration at 10 mg/kg in mice. Data are presented as mean ± s.e.m. of data from three experiments. B: Acute *in vivo* toxicology test was carried out on Balb/c mice. Five control animals received an i.p. injection of saline solution (0.9% NaCl) containing 5% DMSO and five animals received 20 mg/kg of C910 diluted in saline solution with 5% DMSO by the same route. The blood samples were collected after 24 h and then 100 μL of plasma were analyzed by automated biochemical Piccolo Xpress® analyzer (Abaxis). Alanine aminotransferase (ALT), aspartate aminotransferase (AST), amylase (AMY), total bilirubin (TBIL), albumin (ALB), blood urea (BUN), creatinine (CRE), creatine kinase (CK). C-D: Survival (C) and weight (D) of mice following intranasal infection as in Figure 7A with a low dose (5×10^3^ PFU/mouse) of the IAV H1N1 PR/8 strain, and treatment with C910 (*n* = 4) or vehicle control DMSO (*n* = 4). Mantel-Cox test was performed to compare the survival curves between DMSO and C910 groups. Weight loss data are shown as mean ± s.e.m, * *p* < 0.05, multiple *t*-test.

**Supplementary table 1:**
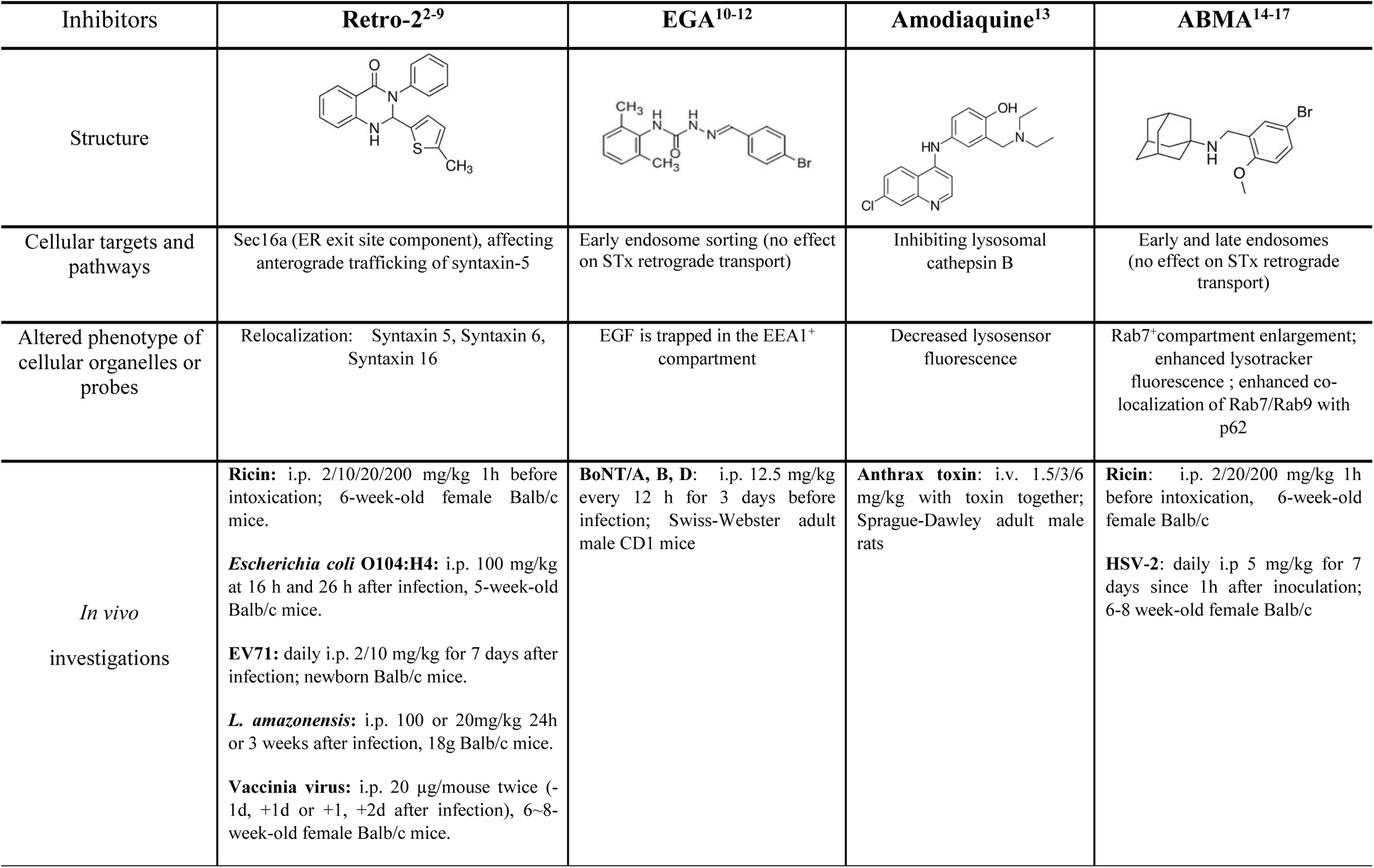
Mode of action of broad-spectrum inhibitors and their *in vivo* investigation.

**Supplementary table 2:**
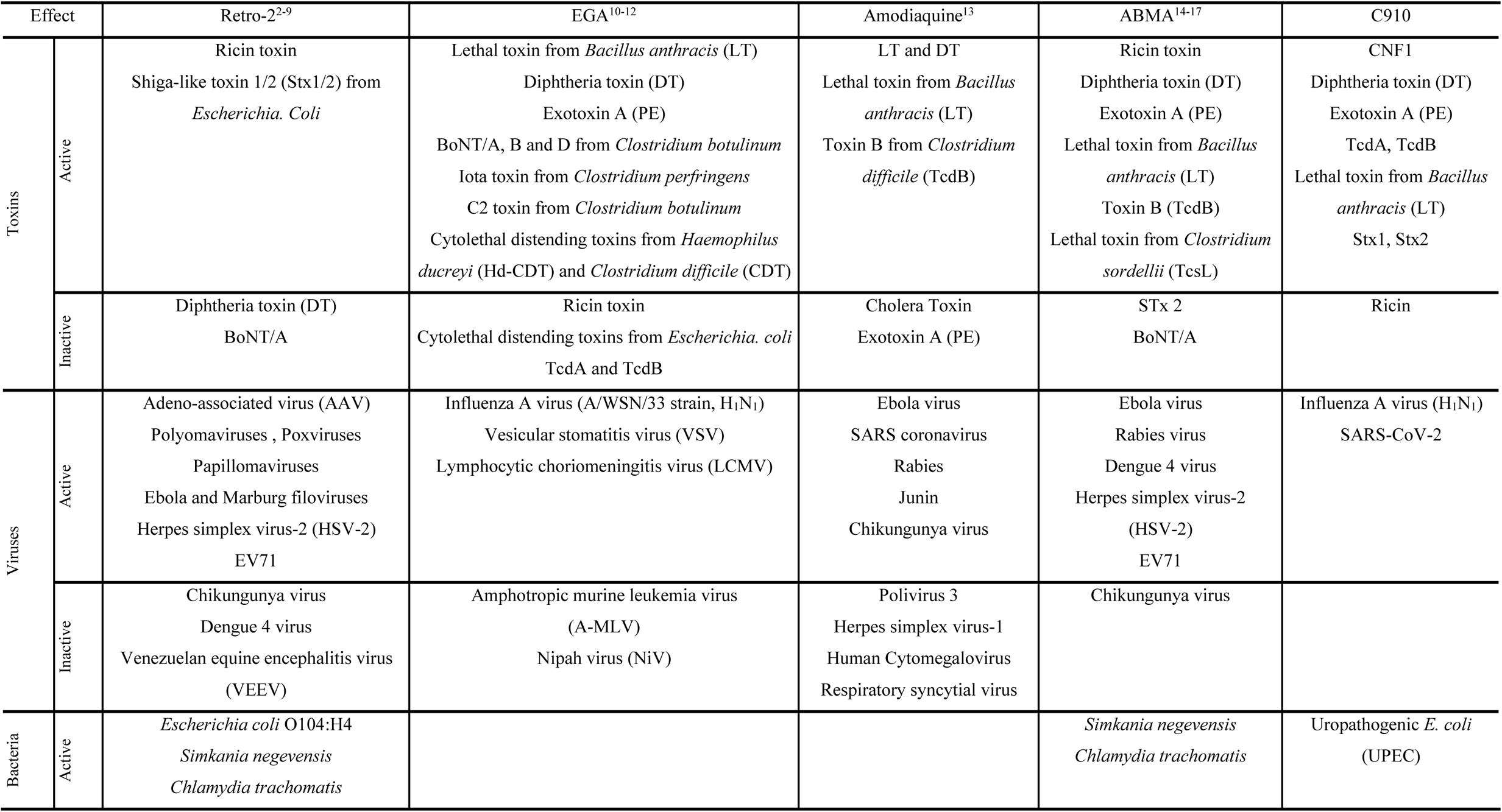
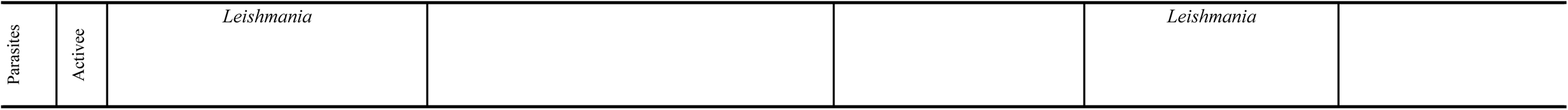
protective spectrum of inhibitors.

## Notes

### Competing Interest Statement

The authors have declared no competing interest.

